# TTLL6-mediated Polyglutamylation of PurA Maintains Colonic Crypt Integrity

**DOI:** 10.64898/2025.12.15.694360

**Authors:** Vanessa Pires, Chloé Pangault, Conception Paul, Pascal Finetti, François Bertucci, Emilie Mamessier, Célia Jardin, Nuttanid Numnoi, Ilaria Cicalini, Damiana Pieragostino, Laura Lebrun, Carsten Janke, Stéphane Audebert, Jean-Paul Borg, Krzysztof Rogowski, Valérie Pinet, Michael Hahne

**Affiliations:** Institut de Génétique Moléculaire de Montpellier, Univ Montpellier, CNRS, Label “Equipe FRM”, Montpellier, France; Cancer Research Center of Marseille (CRCM), Laboratory of Predictive Oncology, Inserm U1068 - CNRS UMR7258 - University of Aix-Marseille UM105 - Paoli Calmettes Institute (IPC), Label “Ligue contre le cancer”, Marseille, France; INSERM, CNRS, Institut Paoli-Calmettes, CRCM, Marseille Protéomique, Aix-Marseille Univ. France; Department of Physiology, Faculty of Medical Science, Naresuan University, Phitsanulok, Thailand; Department of Innovative Technologies in Medicine & Dentistry, Center for Advanced Studies and Technology (CAST), ’G. d’Annunzio’ University of Chieti-Pescara, Chieti, Italy; Institut Curie, Université Paris-Saclay, CNRS UMR3348, 91401 Orsay, France; Tubulin Code team, Institute of Human Genetics, Université Montpellier, CNRS, Label “Equipe FRM”, Montpellier, France

## Abstract

The tubulin tyrosine ligase–like (TTLL) family comprises enzymes catalyzing posttranslational modifications of tubulin, including glutamylation and glycylation. We previously described a critical role for the monoglycylase TTLL3 in colon. Here, we identified TTLL6 as the predominant polyglutamylase in the colon, specifically expressed in epithelial cells of distal and transverse segments. TTLL6 expression decreases during CRC progression, correlating with poor patient prognosis. Deletion of *Ttll6* in mice resulted in elongated colonic crypts, expansion of stem and transit-amplifying compartments, and increased numbers of differentiated epithelial cells. Moreover, *Ttll6*-deficient mice showed an elevated susceptibility to chemically induced colon carcinogenesis. Notably, we identified the nucleic acid–binding protein PurA as a novel TTLL6 substrate with both proteins mutually required for nuclear localization. Consistently, both nuclear polyglutamylation and PurA were present in the bottom compartment of control colons, but reduced in *Ttll6*-deficient colons. These findings reveal a TTLL6-PurA axis being critical in maintaining colonic homeostasis.

## Introduction

Microtubules are the largest filamentous components of the eukaryotic cytoskeleton, consisting of α- and β-tubulin heterodimers that assemble in a polarized manner, with distinct plus and minus ends. Beyond their well-recognized role in conserving cell shape, microtubules are essential in a wide range of cellular processes, such as mitosis, cell motility and differentiation, intracellular signaling and vesicle transport, as well as the formation of specialized organelles. Dysregulation of microtubule dynamics has been implicated in a variety of diseases, ranging from neurological disorders and ciliopathies to cancer.

Posttranslational modifications (PTMs) of tubulin are key regulators of microtubule functional diversity (Magiera and Janke, 2014). These modifications include acetylation and phosphorylation, as well as detyrosination, glycylation and glutamylation. The addition of glycine and glutamate to tubulin are catalyzed by members of the tubulin tyrosine ligase-like (TTLL) enzyme family, in general displaying distinct specificity for initiating or elongating glycine and glutamate side chains, respectively. Both glycylases and glutamylases target the C-terminal tails of tubulin (Janke and Magiera, 2020) and can functionally counterbalance each other (Bosch Grau et al., 2017; Gadadhar et al., 2017). Removal of glutamate side-chains is catalyzed by cytosolic carboxypeptidases (CCPs), also known as deglutamylases, whereas specific deglycylating enzymes have not yet been identified (Janke and Magiera, 2020). More recently also members of Tubulin MetalloCarboxyPeptidases (TMCP) family, were shown to catalyze deglutamylation (Nicot et al., 2023).

We previously demonstrated that *TTLL3* is the only tubulin monoglycylase expressed in colon (Rocha et al., 2014). Tubulin glycylation appears to be restricted to specific primary and motile cilia, as well as flagella (Gadadhar et al., 2017), and indeed *Ttll3*-deficient mice displayed decreased number of primary cilia in the colon (Rocha et al., 2014). While these mice show no overt abnormalities under homeostatic conditions, they display increased susceptibility to chemically induced colon carcinogenesis (Rocha et al., 2014). Furthermore, our analysis of CRC patient biopsies revealed that *TTLL3* transcripts are significantly lower in carcinoma and metastases, as compared to peritumoral tissues, and notably adenoma (Rocha et al., 2014). These findings revealed a role for tubulin glycylation in cancer.

Reports linking TTLL family members with cancer are limited and mostly concern glutamylating enzymes. For example, TTLL11 has been implicated in chromosome mis-segregation and genomic instability (Zadra et al., 2022), while TTLL4 overexpression in breast cancer cells has been associated with enhanced brain metastasis (Arnold et al., 2020). These reports suggest that the role of TTLLs may differ between cancer types, with each enzyme potentially exerting specific functions.

In this study, we investigated the role of tubulin glutamylating enzymes in maintaining colonic homeostasis and their potential impact on tumor development.

## Results

### Ttll6 is the major polyglutamylase in colon epithelium of distal and transverse segments

To determine whether TTLL family members other than the glycylase TTLL3 play a role in colon biology, we assessed glutamylase expression in the colon. qPCR analysis revealed notably high expression levels of the polyglutamylase *Ttll6* specifically in colonic epithelial cells (CECs) (Figure 1A). In contrast, *Ttll6* transcripts were nearly undetectable in the small intestine (Figure 1A). Interestingly, *Ttll6* expression was most prominent in the distal and transverse segments of the colon, as demonstrated by both qPCR and RNAscope (Figures 1B, C).

**Figure 1.**
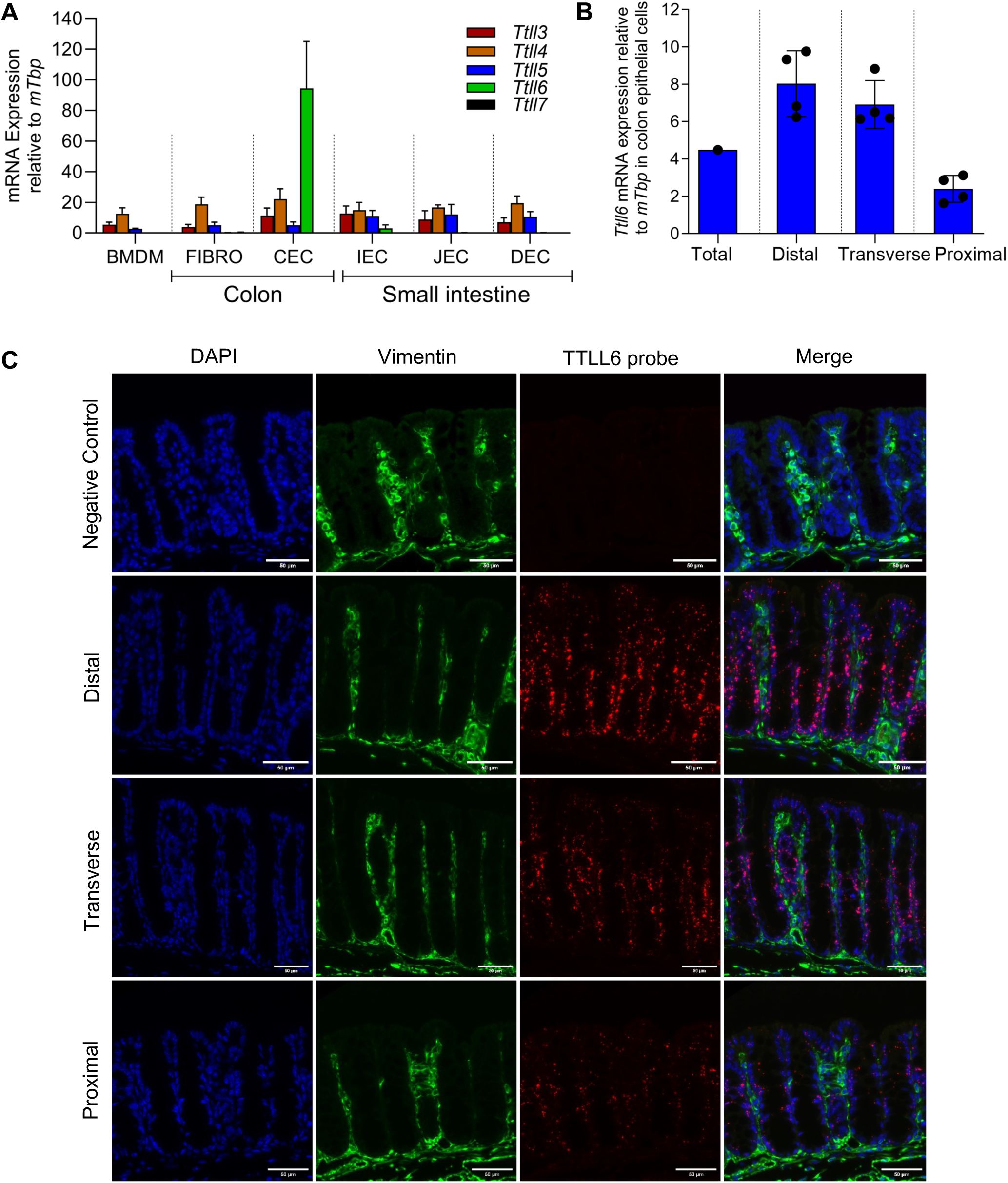
*Ttll6* expression is restricted to epithelial cells in colon. **(A)** qPCR analysis of *Ttll6* expression in different cell population: bone marrow-derived macrophages (BMDM,n=3), colonic fibroblast (FIBRO,n=9), colonic epithelial cells (CEC,n=9), ileal epithelial cells (IEC,n=2), jejunal epithelial cells (JEC,n=2) and duodenal epithelial cells (DEC,n=3). **(B)** qPCR analysis of *Ttll6* expression in colonic epithelial cells (CEC) from the different murine colonic segments (distal, transverse, proximal, n=4). **(C)** *Ttll6* transcript levels in murine tissue sections revealed by RNA scope combined with immunofluorescence. Representative images show in red TTLL6 labelling of transcription foci specifically in CEC with increased intensity in the middle crypt compartment of distal and transverse segments. In green: vimentin-positive fibroblasts in lamina propria; in blue: nuclei stained with DAPI. Scale bar 50µm.

### TTLL6 expression levels decrease during tumor progression in CRC

Our findings in mice prompted us to examine *TTLL6* expression in the CRC patient biopsies we previously reported to exhibit decreased *TTLL3* transcript levels in carcinoma and metastatic tissues as compared to peritumoral regions (Rocha et al., 2014). The analysis displayed a gradual decrease of transcript levels of the glutamylase *TTLL6* in the transition from healthy tissue – adenoma – carcinoma (Figure 2A). Furthermore, analysis of publicly available transcriptome datasets indicated that patients with high *TTLL6*^high^ expression exhibit improved prognosis (Figure 2B). Based on these results, we conducted univariate and multivariate analyses considering clinicopathological parameters and molecular classification. According to the univariate analysis, patient age, sex, tumor location, differentiation grade and microsatellite instability (MSI) status were not significantly associated with recurrence free survival (RFS). In contrast, tumor stage (P=1.39 × 10e-09), the consensus molecular subtypes (CMS) classification (P=4.90 × 10e-03) and *TTLL6*^high^ (P=6.24 × 10e-04) tumors were significantly associated with RFS. Multivariate analysis integrating tumor stage, CMS classification and *TTLL6* data revealed significant association between RFS and tumor stage (P=3.28 × 10e-03 and P=1.42 × 10e-04), but also *TTLL6*^high^ expression (P=4.74 × 10e-02). This is suggesting that *TTLL6*^high^ expression is an independent prognostic factor for RFS (Sup Table 2).

**Figure 2.**
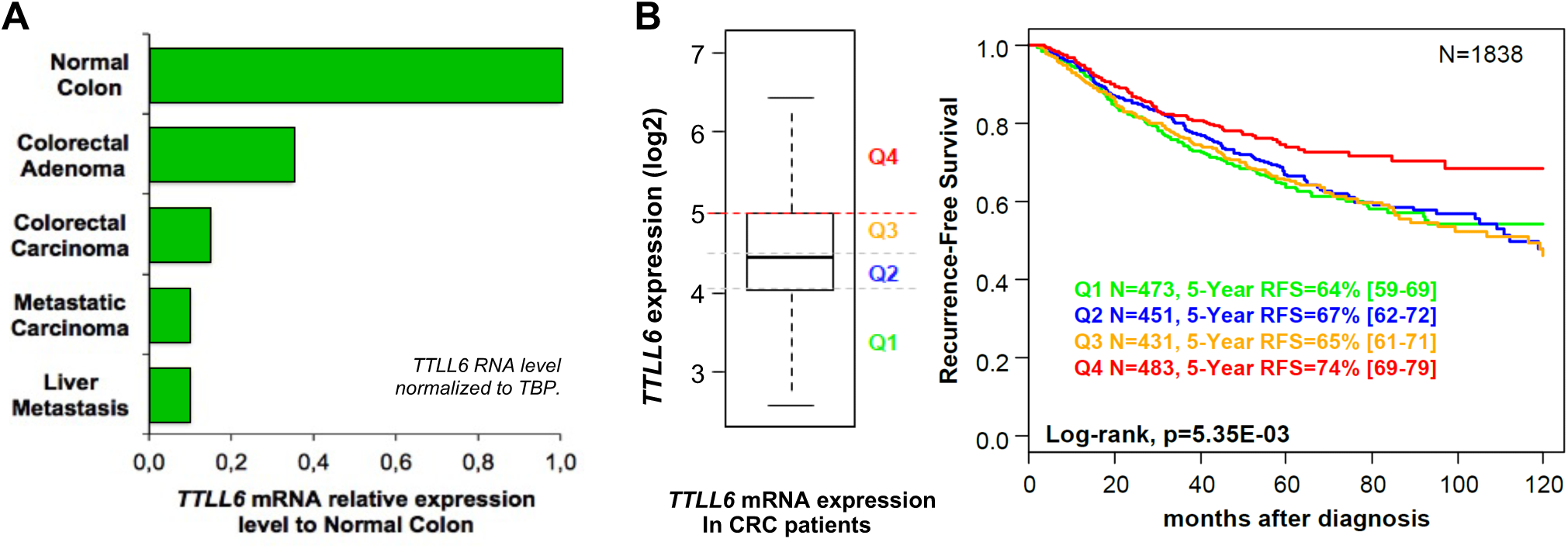
Gradual decrease of *TTLL6* transcript levels during tumour progression. **(A)** Untreated human samples from colorectal adenomas (n=43), primary colorectal carcinomas (n=24), metastatic carcinoma (n=43), liver metastasis (n=27) and matched normal adjacent colorectal tissues (n=101) were analysed for *TTLL6* expression by qPCR. **(B)** High *TTLL6* expressers found in region Q4 (4_th_ quartile) have better Recurrence-Free Survival (RFS). Publicly available transcriptome data of CRC patients (n=1,838) were classified according to *TTLL6* transcript expression levels (left).

### Colons of Ttll6-deficient mice display alterations in different cell compartments

To study the role of TTLL6 in colon biology we established a constitutive *Ttll6 knock in* mouse model (*Ttll6^frt/frt^*, Figure 3A) referred as *Ttll6-*deficient. These mice exhibited no overt phenotypic abnormalities. Based on the prominent expression of *Ttll6* in the distal and transverse colon (Figure 1C), we focused our immunohistological analyses on these segments. We detected a significant increase in crypt length in the transverse colon of *Ttll6*-deficient mice (Figure 3B), which correlated with an increased number of cells per crypt (Figure 3C). To examine cell proliferation, we performed immunostaining for Ki-67, a marker of actively dividing cells. Crypts were divided into 3 compartments, as illustrated in figure 3D. In the transverse segments of *Ttll6*-deficient colons, both the bottom and middle crypt compartments exhibited a marked increase in Ki-67–positive cells (Figure 3E). Moreover, transverse compartments showed elevated cell number in S-phase of cell cycle, as determined by BrdU incorporation assays (Figure 3F). The middle compartments of distal crypts from *Ttll6*-deficient colons also displayed elevated Ki-67 staining, but no increased levels of BrdU incorporation (Supplemental Figure S1).

**Figure 3.**
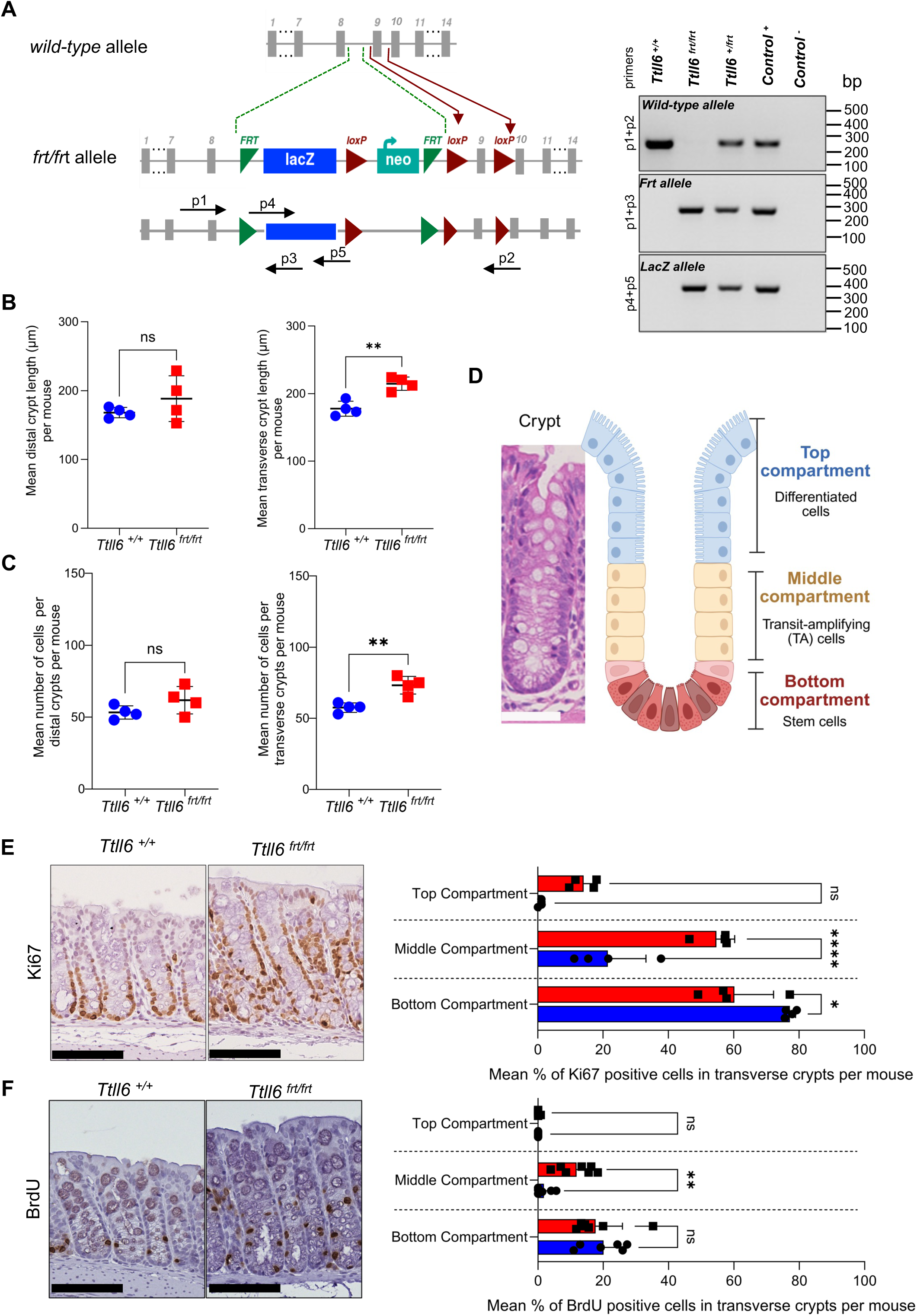
*Ttll6*-deficient mice show increased crypt length and cell proliferation. **(A)** Generation of *Ttll6 knock in* mice (*Ttll6 _frt/frt_*). A gene-trap cassette was inserted into the *Ttll6* gene (top schematic) between exons 8 and 10 by homologous recombination in mouse embryonic stem cells to generate a “knockout-first” allele (frt). The cassette includes a reporter gene (*LacZ* in blue) and a selectable marker (neomycin(neo) in light blue) and exons are indicated by grey boxes. The cassette contains flippase (frt, green) and Cre (loxP, dark red) recombinase recognition sites. Representation of primers (black arrows) used for genotyping. Genomic DNA was extracted from mice tails and PCR performed using primer 1 (p1) and primer 2(p2) (product 259 base pair (bp) to detect the wild-type allele; primer 1 and primer 3 (p3)(product 248bp) to identify the *frt* allele; in addition primer 4 (p4) and primer 5 (p5) (product 400bp) were employed to reveal *LacZ*. **(B)** Measurement of crypt length in distal and transverse segments (n=4 for each group *Ttll6_+/+_* (blue) and *Ttll6_frt/frt_* (red)). **(C)** Respective cell count of the crypts measured in B. **(D)** Schematic representation of an open colonic crypt in homeostasis in mouse. Colonic crypt are composed by different types of epithelial cells and can be divided into three compartments: stem cells (bottom; red), transit amplifying (TA) cells (middle; yellow) and differentiated cells (top; blue). **(E)** Representative images of immunohistochemical analysis for Ki67 in transverse segments (left) and respective quantification as percentage of Ki67-positive cells per crypt compartment (right; bars in blue and red represent values for *Ttll6_+/+_* and *Ttll6_frt/frt_* animals, respectively). Dots represent mean values of crypts analysed per mouse (n=4 for each group). **(F)** Representative images of transverse colon sections from mice following a 2-hour BrdU pulse (left), stained for BrdU incorporation (n=6 for *Ttll6_+/+_* (blue) and n=7 for *Ttll6_frt/frt_* mice (red)) with the respective quantification per crypt compartments (shown as % (right). Dots represent mean values of crypts analysed per mouse. Per mouse 10-40 open crypts were quantified per segment (B,C,E,F). Scale bar 100µm. Statistical analysis in B, C, E,F: ns p>0.05, * p < 0.05, **p < 0.01, ***p < 0.001, ****p < 10−4 by two-tailed unpaired t-test (B,C) and ordinary two way ANOVA-Šídák’s multiple comparisons test (E,F).

Next, we analyzed potential physiological alterations in the different CEC populations. To examine the stem/progenitor cell compartment, we performed immunostaining for CD44v6, a known marker of colonic crypt base (Afify et al., 2016; Nagata et al., 2019). Indeed, CD44v6 expression was confined to the base of the colonic crypts in control mice, but expanded along the crypt axis *Ttll6*-deficient colons (Figure 4A). We next assessed the presence of differentiated epithelial cell types. Alcian blue staining revealed an increased number of mucin-producing cells in the distal segment of *Ttll6*-deficient colons, which was also detectable by Mucin 2 immunostaining (Figures 4B, Supplemental Figure S2A, B). In contrast, the number of enteroendocrine and tuft cells, determined by immunostaining for chromogranin A and Dclk1 respectively, remained unchanged (Supplemental Figure S2C). qPCR analysis showed elevated expression of *Retnlb*, a goblet cell derived protein, and *Alpi*, which is expressed by enterocytes (Jones and Dempsey, 2016), while transcripts of *Chga*, the gene encoding chromogranin A, and *Dclk1* remained unchanged (Figure 4C,D).Taken together, these findings indicate that *Ttll6* contributes to the maintenance of colonic crypt integrity.

**Figure 4.**
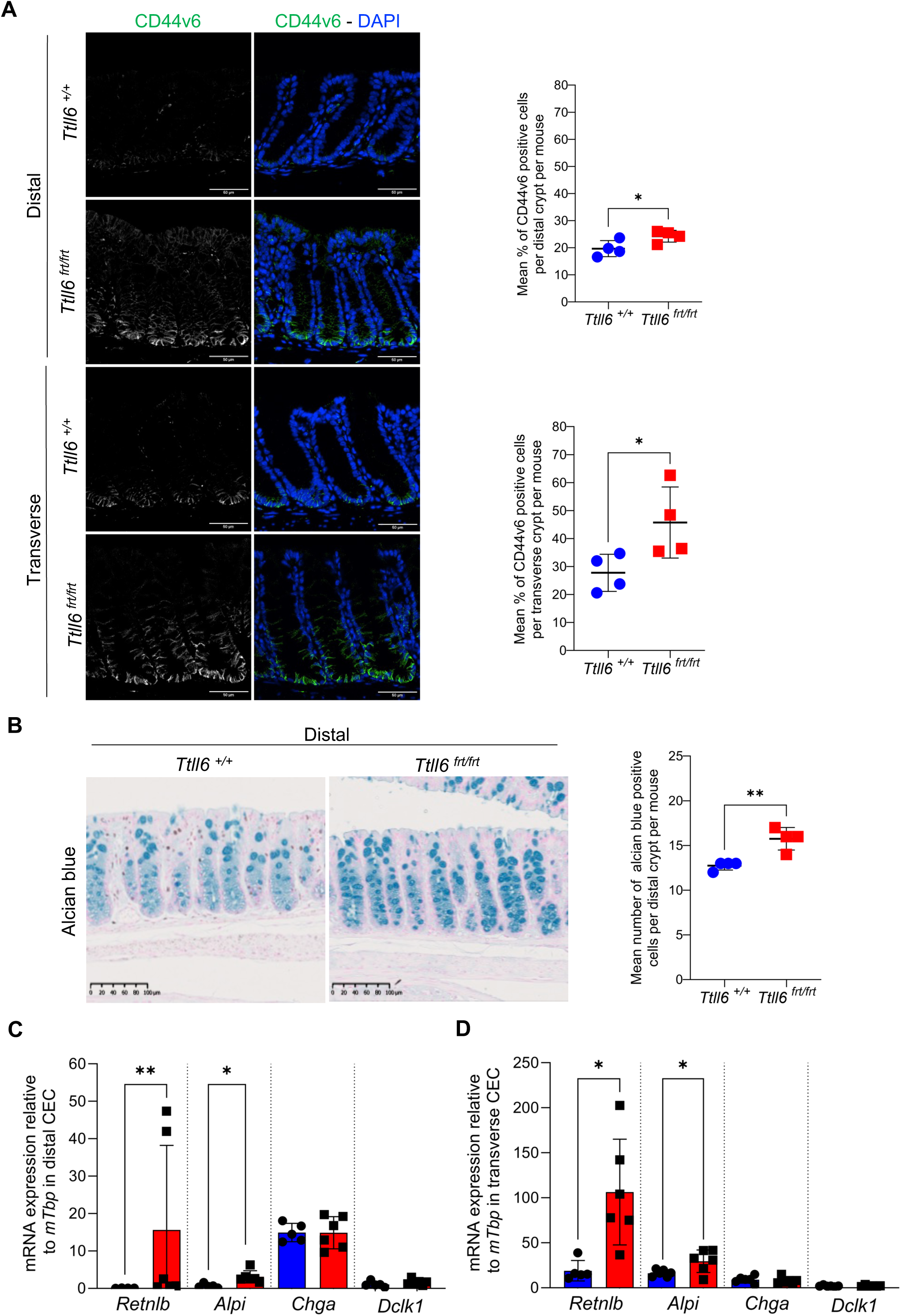
*Ttll6*-deficient mice exhibit disruptions in colonic crypt organization and cell populations. **(A)** Representative images of immunostaining for the stem compartment marker (CD44v6) in distal and transverse segments. The right panel shows the respective quantification (n=4 for each group *Ttll6_+/+_* (blue) and *Ttll6_frt/frt_* (red)). Scale Bar 50µm. **(B)** Representative images of immunohistochemical analysis for mucin positive cells in distal segments, visualized using Alcian blue and respective quantification of absolute number alcian blue positive cells per crypt per mouse in distal segment (n=4 for each group *Ttll6_+/+_* (blue) and *Ttll6_frt/frt_* (red)). Scale bar 100µm **(C)** qPCR analysis of known markers of differentiated cells, as *Retnlb* (Deep secretory cells, n= 4 for *Ttll6_+/+_* (blue),n=6 for *Ttll6_frt/frt_* (red)), *Alpi* (enterocyte progenitors / enterocyte cells, n=5 for *Ttll6_+/+_* (blue), n=6 for *Ttll6_frt/frt_* (red)), *Chga* (ChromograninA, enteroenendocrine cells, n=5 for *Ttll6_+/+_* (blue), n=6 for *Ttll6_frt/frt_* (red)) and Dclk1(Tuft cells, n=5 for *Ttll6_+/+_* (blue), n=6 for *Ttll6_frt/frt_* (red)) expression in colonic epithelial cells (CEC) from distal segment.**(D)** qPCR analysis of known markers of differentiated cells, as *Retnlb* (Deep secretory cells, n= 5 for *Ttll6_+/+_* (blue),n=6 for *Ttll6_frt/frt_* (red)), *Alpi* (enterocyte progenitors / enterocyte cells, n=6 for *Ttll6_+/+_* (blue), n=6 for *Ttll6_frt/frt_* (red)), *Chga* (ChromograninA, enteroenendocrine cells, n=6 for *Ttll6_+/+_* (blue), n=6 for *Ttll6_frt/frt_* (red)) and Dclk1(Tuft cells, n=6 for *Ttll6_+/+_* (blue), n=6 for *Ttll6_frt/frt_* (red)) expression in colonic epithelial cells (CEC) from transverse segment Per mouse 10-40 open crypts were quantified per segment. Statistical analysis : ns p>0.05, * p < 0.05, **p < 0.01, ***p < 0.001, ****p < 10−4 by two-tailed unpaired t-test.

### Ttll6-deletion in colonic epithelial cells modulates colon carcinogenesis

We then wondered whether *Ttll6*-deficiency in CEC alters the susceptibility to colon carcinogenesis. To address this, we generated a *Ttll6^flx/flx^ villin-creErt* mouse strain, in which *Ttll6* can be specifically deleted in the epithelial compartment of colon upon tamoxifen administration. Efficiency of tamoxifen induced *Ttll6*-deletion in CEC was validated by qPCR (Supplemental Figure S3). To study colon carcinogenesis we employed a protocol of chemically induced colitis-associated colon carcinogenesis (CAC; Figure 5A), which mimics the sequential development of human colon carcinogenesis i.e., the crypt-foci-adenoma-carcinoma sequence (Kucherlapati, 2023; Tanaka, 2012).

**Figure 5.**
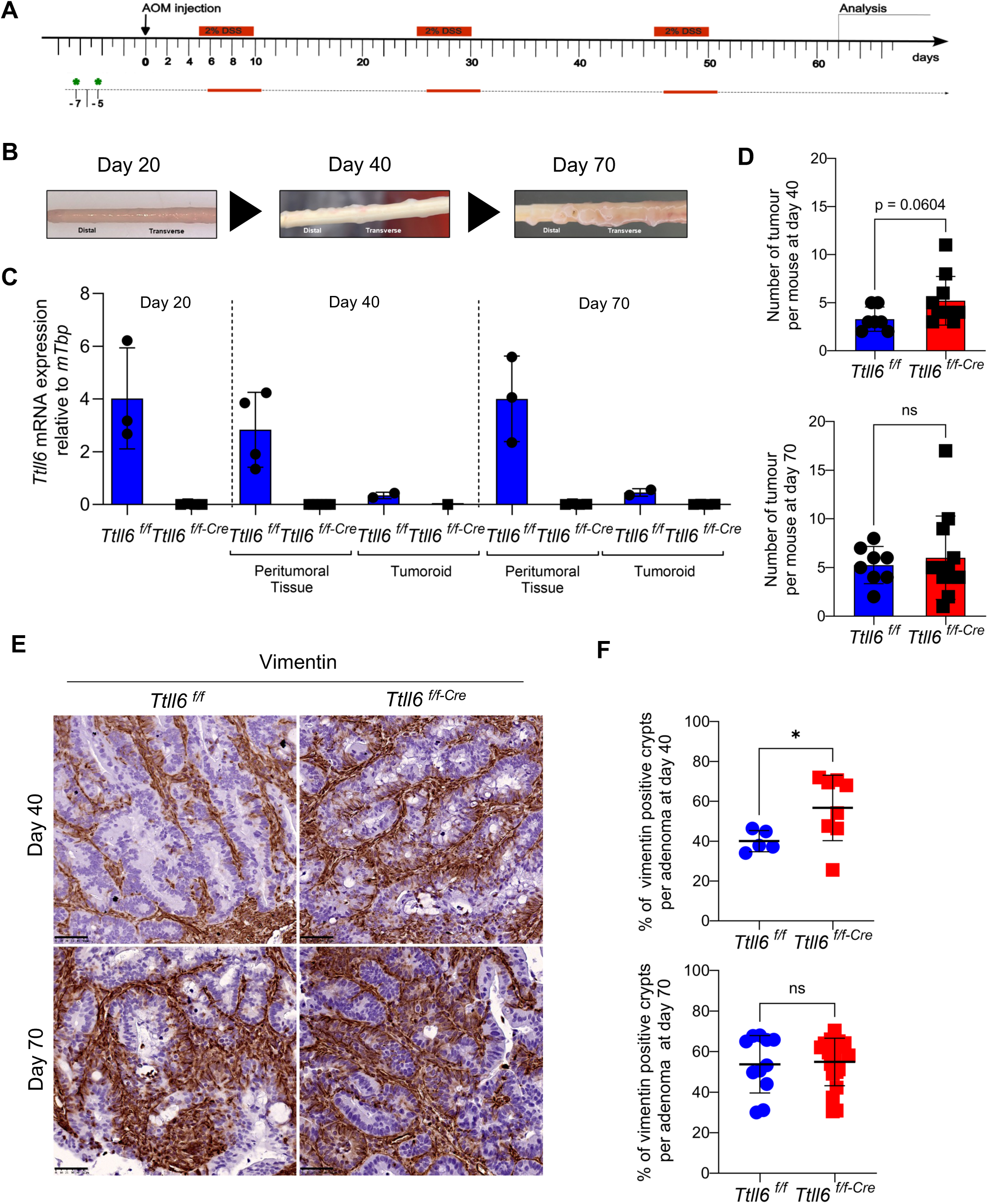
Impact of *Ttll6*-deficiency in colon carcinogenesis. **(A)** Schematic representation of the colitis-associated carcinogenesis (CAC) protocol. *Ttll6* was deleted in *Ttll6_f/f-Cre_* mice by two consecutive tamoxifen injections before starting the protocol (green asterisk). **(B)** Representative images of colons collected on day 20, day 40 and day 70, respectively. **(C)** *Ttll6* transcript levels determined by qPCR during carcinogenesis in wild-type (*Ttll6_f/f_*) and *Ttll6*-deficient (*Ttll6_f/f-Cre_*) mice of colon epithelial cells from distal and transverse segments at day 20, and peritumoral tissue (n=3) and tumoroids (n=2) obtained from samples isolated at days 40 and 70. *Ttll6* transcript levels were normalized to mouse TATA-binding protein (*Tbp*). **(D)** Number of tumoral lesions per mouse in colons collected at day 40 and day 70 in wild-type (*Ttll6_f/f_*; blue bar), and Ttll6-deficient (*Ttll6_f/f-Cre_*; red bar). **(E)** Representative images of immunohistochemistry for vimentin in adenomas at day 40 and day 70. Scale bar 50µm. (F) Respective quantification of vimentin positive crypts in adenomas at day 40 (n=6 for *Ttll6_f/f_* (blue) and n=8 for *Ttll6_f/f-Cre_* (red) animals, and day 70 (n=11 for *Ttll6_f/f_*, n=20 for *Ttll6_f/f-Cre_*, n represents number of adenoma analysed) Statistical analysis in D,F : ns p>0.05, * p < 0.05, **p < 0.01, ***p < 0.001, ****p < 10−4 by two-tailed unpaired

To monitor *Ttll6* levels in CEC and tumors during the CAC protocol (Figure 5A), mice received two tamoxifen injections prior to protocol initiation, and colons were collected at days 20, 40, and 70 (Figure 5B). Macroscopic tumor lesions were detectable only at days 40 and 70, allowing their isolation. qPCR analysis confirmed that deletion of *Ttll6* was maintained in *Ttll6^flx/flx^ villin-creErt* mice throughout the CAC protocol (Figure 5C). Notably, tumor lesions in control mice revealed significant downregulation of *Ttll6* in tumor lesions, whereas the surrounding areas maintained high *Ttll6* levels (Figure 5C). Strikingly, the numbers of macroscopic lesions were elevated at day 40 in *Ttll6^flx/flx^ villin-creErt* mice showing a trend toward significance, but were comparable in *Ttll6^flx/flx^ villin-creErt* and control animals at day 70 (Figure 5D). Size of tumor lesions were similar in colons of control and *Ttll6*-deficient animals both at days 40 and 70. Importantly, when we scored for vimentin-positive crypts, a marker of tumor aggressiveness (Ma et al., 2019), the tumors in *Ttll6*-deficient colons exhibited an increased incidence of this type of crypts (Figures 5 E,F).

Together, in the context of CAC, *Ttll6* deletion appears to promote early tumor development.

### Polyglutamylation is not detectable in primary cilia of colonic epithelial cells

We previously reported that, in the colon, primary cilia (PC) are predominantly found on fibroblasts and are rarely observed on epithelial cells (Paul et al., 2022). Reports describe an important role for TTLL6 in the maintenance of PC stability (He et al., 2018; Pathak et al., 2011). We therefore analyzed the polyglutamylation status of PC in colon using the polyE antibody, which recognizes C-terminally exposed amino acid chains composed of at least 3 glutamates (Rogowski et al., 2010). As shown in Figure 6A, polyglutamylation was absent in PC of epithelial cells but present in those of colonic fibroblasts. Notably, polyE immunostaining was detectable in nuclei of CEC, which was decreased in CEC of *Ttll6*-deficient mice (Figure 6B). Nuclei of fibroblasts showed no immunostaining for polyE. In agreement with the TTLL6 expression patterns depicted in Figure 1, polyglutamylation levels in the colon exhibited a gradient, highest in the distal segment and diminishing toward the proximal segment (Sup. Figure 4). These findings indicate that polyglutamylation might play distinct cell-type specific function. In fibroblast it contributes to the stabilization of PC through the modification of tubulin, while in epithelial cells it has a nucleus-associated function.

**Figure 6.**
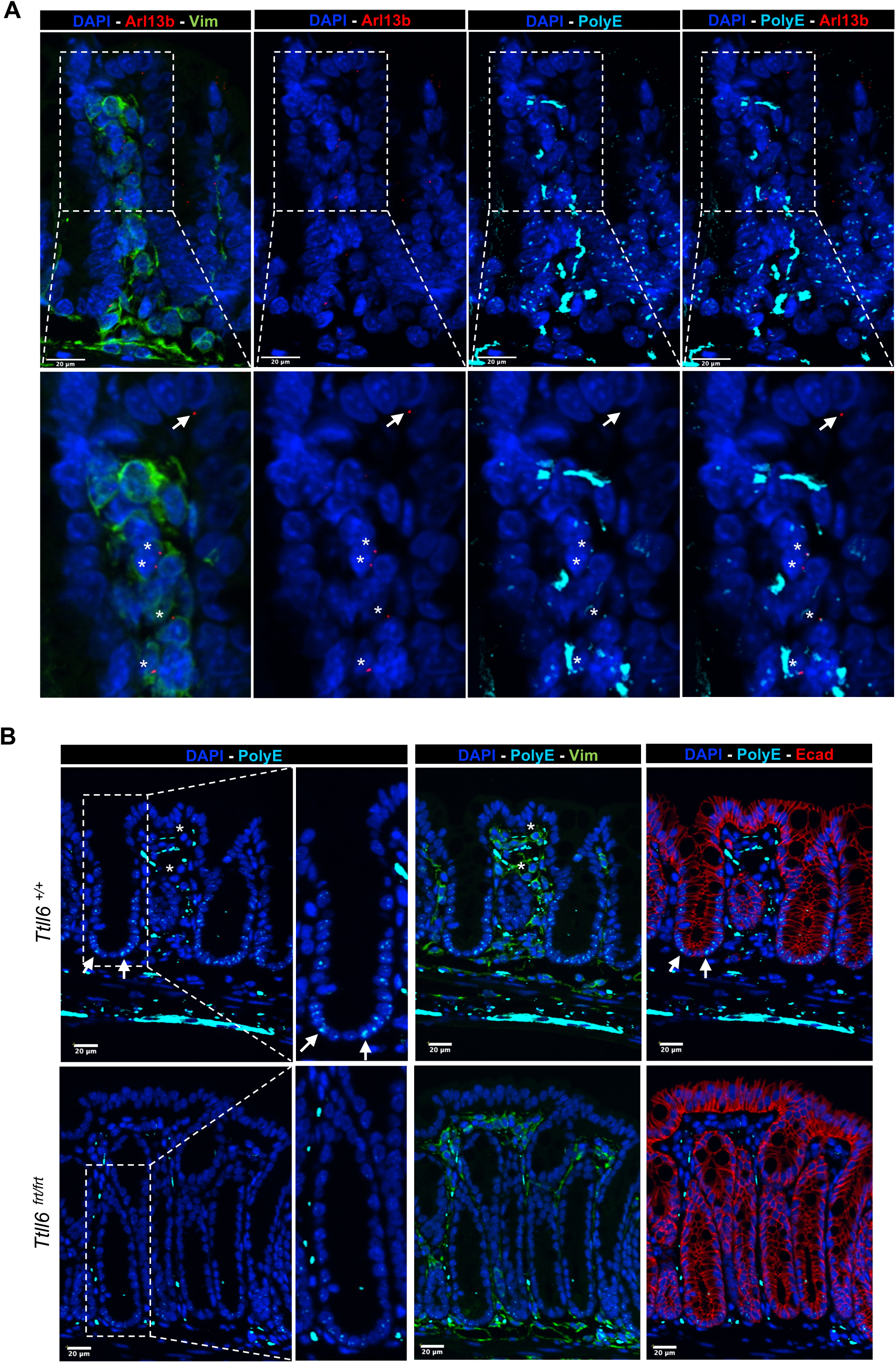
Polyglutamylation status is different between colonic fibroblasts and epithelial cells. **(A)** Polyglutamylation (in cyan) is not detectable in primary cilia (in red) of colonic epithelial cells. Primary cilia labeled by Arl13b immunostaining in red. Arrows label epithelial cells and asterisks (*) fibroblasts, identified by vimentin staining (in green). Lower panel shows blow-up of corresponding upper panel images. **(B)** Nuclei of colonic epithelial cells, but not fibroblasts display polyglutamylation. Upper panel: In distal colons of control mice (*Ttll6_+/+_)*, epithelial cells labeled with E-Cadherin (in red and highlighted by arrows) exhibited a nuclear punctuated staining for polyglutamylation (in cyan), but not fibroblasts labeled with vimentin (in green and marked by asterisks (*)). Lower panel: In *distal* colon of *Ttll6_frt/frt_ mice,* polyglutamylation levels are reduced in nuclei of epithelial cell nuclei compared with those of *Ttll6_+/+_* animals. Nuclei were stained with DAPI (in blue). Scale bars represent 20µm. Magnifications of the DAPI-PolyE staining are two-fold.

### Identification of candidate substrates of TTLL6 in CEC

Although tubulin is the canonical substrate for glutamylation, a growing number of additional proteins have been suggested to undergo this modification (van Dijk et al., 2008; Kashiwaya et al., 2010; Ruse et al., 2022; Kravec et al., 2024). Our observation that polyglutamylation is detectable in the nuclei of CEC, which under normal conditions are devoid of tubulin, suggests that TTLL6 is catalyzing polyglutamylation of alternative substrates. To identify novel glutamylated proteins in CECs, we performed immunoprecipitation from wild-type CEC lysates using the polyE antibody. Immunoprecipitates were compared with pull-downs performed using plain beads lacking antibody to assess specificity. Recovered proteins were subjected to mass-spectrometry–based proteomic analysis, revealing ten proteins significantly enriched in the polyE IP condition (Sup. Table 3), including two RNA/DNA binding proteins, PurA and PurB (purine-rich element binding protein A/B) known to interact with each other (Figure 7A left panel). Comparison of immunoprecipitates from wild-type and *Ttll6*-deficient CEC identified 43 enriched proteins including several putative RNA-binding proteins (RBPs) (Sup. Table 4; Figure 7A right panel). Among the identified candidates, PurA and PurB were of particular interest as their recovery by PolyE immunoprecipitation was lower in *Ttll6*-deficient CECs. In addition, PurA contains a C-terminal tail, which bears a strikingly resemblance with alpha-tubulin tail (Figure 7B).

**Figure 7.**
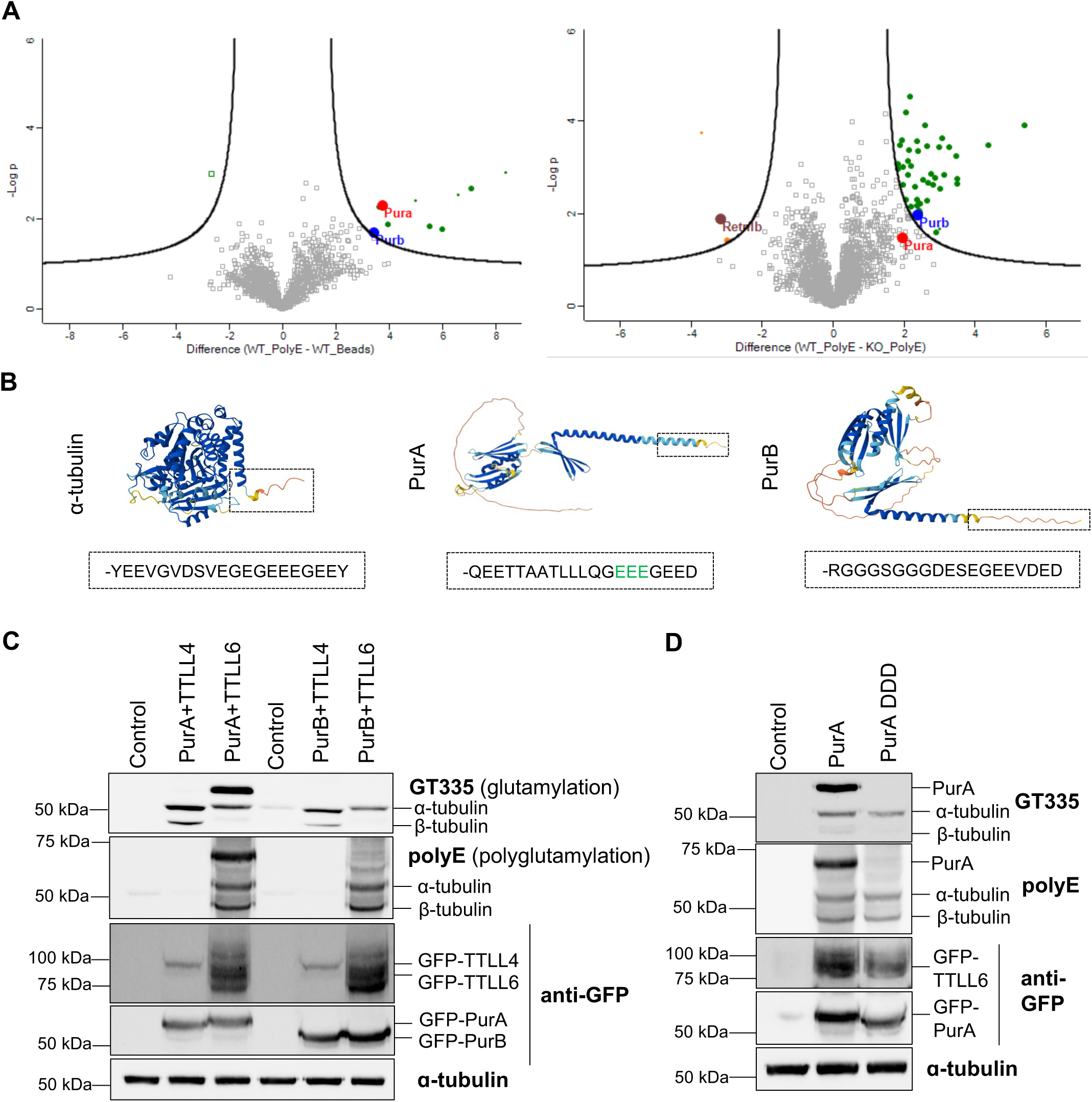
*PURA is a novel substrate of TTLL6*. **(A)** Volcano plots showing differential protein enrichment identified by mass spectrometry following immunoprecipitation of polyglutamylated proteins from colonic epithelial cells. Left panel: Comparison between immunoprecipitates from *Ttll6_+/+_* (WT_PolyE) CEC lysates using the polyE antibody and pull-downs performed with beads lacking antibody (WT_Beads). Proteins above black line are statistically different between the two experimental groups and highlighted, including PurA (red dot) and PurB (blue dot). Right panel: Comparison between polyE immunoprecipitates from wild-type Ttll6_+/+_ (WT_PolyE) and Ttll6_frt/frt_ (KO_PolyE) CEC lysates displaying decreased recovery in *Ttll6_frt/frt_* lysates, while Retnlb (brown dot) is increased. **(B)** Alpha-fold protein structure prediction of α-tubulin, PurA and PurB. In the dashed black box, the C-terminal region of each protein with the respective last twenty amino acid sequence, displaying high similarity in the last eight amino acids of PurA and α-tubulin. **(C)** Immunoblot of protein extracts from HEK293 cells co-expressing PurA or PurB full length (-EGFP tag) together with TTLL4-EGFP or TTLL6-EGFP, as indicated. Notably, glutamylation (GT335) and polyglutamyation (polyE) as revealed by GT335 and polyE antibodies, respectively, are only present in cells co-expressing PurA with TTLL6-EGFP and not TTLL4-EGFP. No glutamylation of PurB is detectable. **(D)** Immunoblot of protein extracts from HEK293 cells co-transfected with TTLL6-EGFP together with either full-length PurA (PurA-EGFP) or mutated PurA DDD (PurA DDD-EGFP, glutamate residues (E) replaced by aspartate (D), highlighted in green in (B)).

To determine whether PurA or PurB are substrates of TTLL6, we co-transfected human embryonic kidney (HEK) cells with PurA or PurB together with either TTLL6 or TTLL4, a monoglutamylase known to modify numerous non-tubulin substrates (van Dijk et al., 2008). Strikingly, TTLL6 but not TTLL4 catalyzed glutamylation of PurA, whereas neither enzyme modified PurB (Figure 7C). To confirm that glutamate residues in the C-terminal stretch of PurA are indeed the target TTLL6-dependent modification, we generated a mutated version of PurA by replacing the 3 glutamates present within the C-terminus of PURA with aspartates, which can no-longer be modified (PurA-DDD) (Figure 7B). Indeed, in cells co-transfected with PurA-DDD and TTLL6, the polyE antibody failed to detect any polyglutamylation signal (Figure 7D).

PurA (PURA) can be localized in different cellular compartments including cytoplasm and nucleus (Gallia et al., 2000), which prompted us to test whether the cellular location of PurA is altered by its glutamylation status. For this, retinal pigment epithelial (RPE) cells were co-transfected with TTLL6-EGFP and PurA-HA (full-length) or the variant PurA-DDD-HA. As illustrated in Figure 8A and Sup. Figure 6B, PurA was primarily localized in the nucleus, whereas PurA-DDD was mostly detected in the cytoplasm. Remarkably, TTLL6 was prominently detected in the nucleus of PurA-transfected cells, whereas its nuclear presence was markedly reduced in PurA-DDD transfectants.

**Figure 8.**
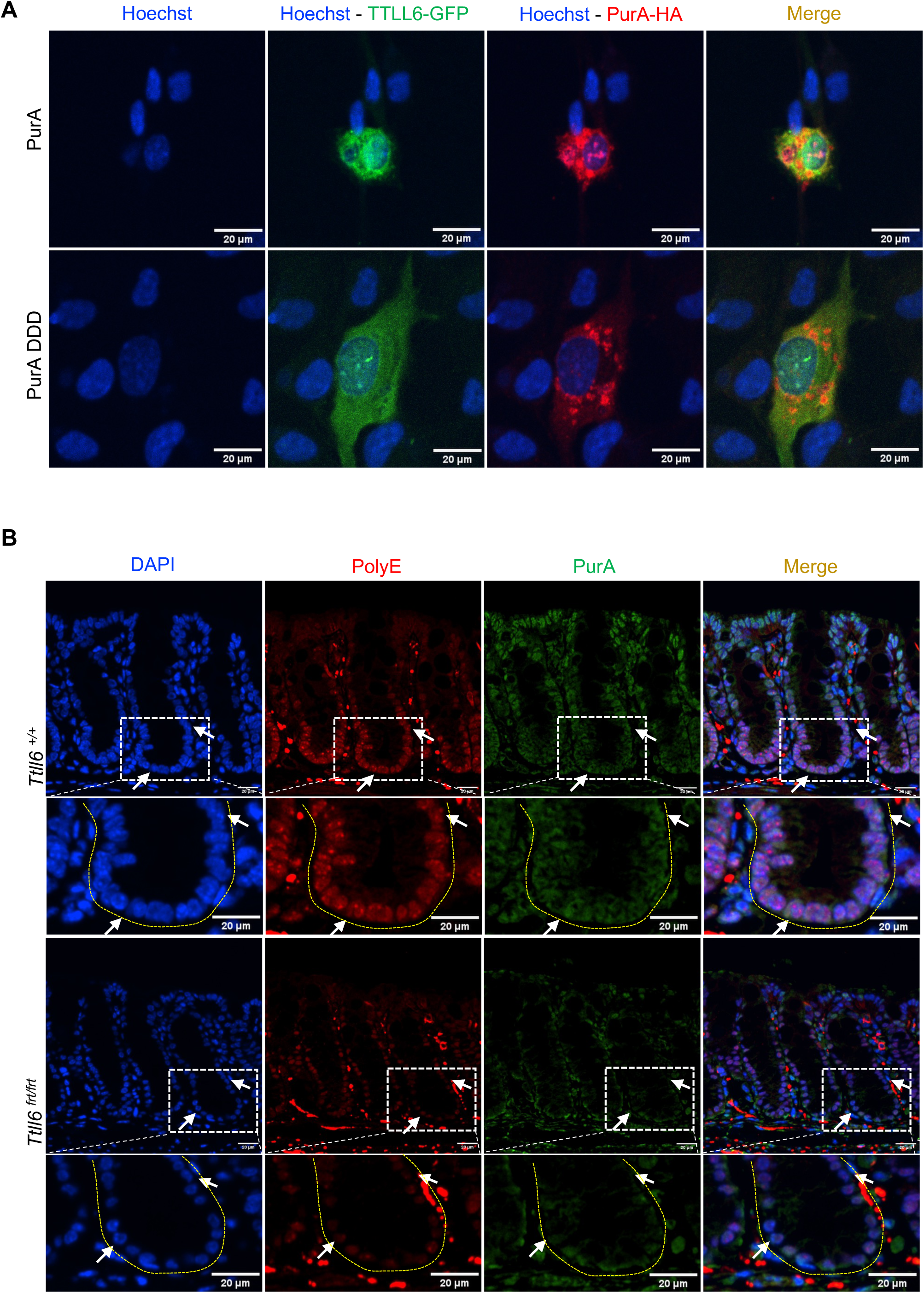
PurA accumulates in the nucleus in presence of *Ttll6*. **(A)** Representative images of immunofluorescence analysis of retinal pigment epithelial (RPE) cells co-transfected with TTLL6-EGFP and PurA-HA (full-length) or PurA-DDD-HA (mutated form). PurA (in red) and TTLL6 (in green) are detectable in nucleus as well as cytoplasm (top panel). In contrast, in cells co-transfected with TTLL6 and PurA mutated (PurA DDD), only TTLL6 is detected in nucleus and cytoplasm, whereas PurA DDD is located mainly in the cytoplasm (bottom panel). **(B)** Representative images of immunostaining on mouse colon for polyglutamylation (polyE in red) and purA (in green). In *Ttll6 _+/+_* mice, polyE staining is detectable in bottom and middle crypt compartments with accumulation in nuclear foci (arrow). PurA is present in all nuclei of CEC in the crypt. In *Ttll6 _frt/frt_* mice, polyE staining is reduced and not detectable in nuclear foci; purA stainig in nuclei is reduced in bottom and middle crypt compartment (arrow), and in the top compartment not all nuclei reveal PurA staining. Additional images are shown in supplementary figure 8 collected from 3 control and 3 *Ttll6*-deficient mice. Scale bar 20um.

To test whether glutamylation-dependent nuclear re-localization of PURA observed *in cellulo* also occurs in colon tissue, we employed an antibody that specifically recognizes PurA, but not PurB (Supplementary Figure 6A and Figure 8), enabling the assessment of PurA expression in the colon of control and *Ttll6*-deficient mice. Nuclear staining of PurA in CEC was markedly reduced in the bottom and middle compartments of the crypts in *Ttll6*-deficient tissues (Figure 8B and Sup. Figure 7). Our data indicate that PurA is a novel substrate of TTLL6 polyglutamylase and its cellular localization appears to be modulated by its glutamylation status.

## Discussion

TTLL6 is a member of the tubulin tyrosine ligase-like (TTLL) enzyme family known to add glutamate at the C-terminal tails of alpha-tubulin (van Dijk et al., 2007). Since its initial identification, the function importance of TTLL6 apart from its role in cilia remains poorly understood. Several reports have associated TTLL6 with the regulation of microtubules in the axoneme of cilia as well as neurons (Bosch Grau et al., 2013; Pathak et al., 2011; Zempel et al., 2013). Our study is the first to describe a role for TTLL6 in colon, where its expression is restricted specifically to the epithelial compartment.

We previously reported that CEC exhibit a relatively low abundance of primary cilia (Paul et al., 2022), which contrasts with epithelial cells in other tissues, such as the kidney and respiratory tract. This is likely attributable to the rapid turnover of intestinal epithelial cells, occurring approximately every five days (Maurizy et al., 2021). Actively proliferating cells generally do not form primary cilia, as centrioles are engaged in mitotic spindle formation and unavailable for basal body assembly (Atmakuru and Dhawan, 2023). Accordingly, we previously described that in colon PC are rarely present on epithelial cells but are mostly present in fibroblast (Paul et al., 2022). Notably, Immunostaining for polyE on colon sections revealed that PC in epithelial cells are not polyglutamylated, in contrast to PC in fibroblasts, which suggests a distinct composition and possibly specific function of PC in the different cellular compartments of colon. Indeed, it has been recently reported that a large part of ciliary proteomes is cell type specific (Hansen et al., 2025).

A more recent report linked TTLL6 with cancer biology. Zhong and colleagues described that the transcription factor XBP1v1 promotes TTLL6 expression thereby triggering mitotic spindle polyglutamylation and that TTLL6 knockdown in a human breast cancer cell line induces cell death (Zhong et al., 2022). In contrast, we observed a downregulation of *TTLL6* transcript levels during disease progression in biopsies of CRC patients. Consistently, transcriptome datasets suggest that CRC patients with low TTLL6 expression levels have a poorer prognosis. Together, these findings point to cancer type specific functions of TTLL6. Interestingly, our data suggest that TTLL6 downregulation contributes early to the development of preneoplastic lesions, in contrast to TTLL3, whose transcription is decreased at later stage during progression to malignancy (Rocha et al., 2014). Together, both glycylation and glutamylation appear to play a distinct role in colorectal carcinogenesis.

We found that *Ttll6* expression is predominantly localized to epithelial cells in the distal and transverse segments of the colon, regions known to be more susceptible to tumorigenesis in the CAC model (Felton et al., 2018). Consistently, the most pronounced changes in *Ttll6*-deficient colons were observed in the transverse region, where crypts were markedly elongated. This elongation corresponds to an expansion of specific functional units within the crypt, particularly the stem cell and transit-amplifying (TA) compartments, as indicated by CD44v6 and Ki-67 immunolabeling. Moreover, there was an increase in cells expressing differentiated markers, including mucin- and Retnlb1-positive secretory cells, such as goblet cells, and Alpi-positive enterocytes. The observed changes in these regions, consistent with the restricted expression of *Ttll6*, likely reflect an expansion or reorganization of the stem and progenitor (TA) compartments in its absence.

It is known that non-tubulin proteins can also be glutamylated. Well-characterized examples are the histone chaperones nucleosome-assembly proteins or members of the nucleophosmin / nucleoplasmin family (van Dijk et al., 2008). The modification of non-tubulin substrates have so far been mostly shown to be catalyzed by glutamylases distinct from TTLL6, though Xia and colleagues reported that TTLL6 specifically regulates the polyglutamylation of cGAS, thereby modulating type I interferon responses (Xia et al., 2016). In this work we identified a novel substrate for TTLL6, PurA, a nucleic acid binding protein. While on tubulin mammalian TTLL6 was shown to be a predominant glutamate chain elongating enzyme, it can also initiate glutamate chains (van Dijk et al., 2007). This is consistent with our finding that TTLL6 alone is sufficient to establish polyglutamylation at the C-terminal tail of PurA, as identified using the polyE antibody, which specifically recognizes chains composed of at least three glutamates. The C-terminal tail of PurA closely resembles that of tubulins, the canonical substrates of TTLL6, in particular the stretch of three glutamate residues that were validated as polyglutamylation sites by site-directed mutagenesis.

Co-transfection experiments identified a mutual dependency of TTLL6 catalyzed polyglutamylation of PurA and the localization of both proteins. Indeed, disruption of the glutamate stretch in the C-terminus of PurA restricted the localization of the mutated variant, as well as TTLL6 to the cytosol. Interestingly, in tissue, polyE immunostaining revealed polyglutamylation particularly in the nuclei of the epithelial stem cell compartment in distal and transverse segments of colon, which was significantly reduced in *Ttll6*-deficient colons.

Few studies have reported an oncogenic role for PurA. Loss of nuclear PurA has been linked to androgen receptor overexpression and tumor progression in prostate cancer, whereas in esophageal squamous cell carcinoma, PurA promotes metastasis through activation of mesoderm-specific transcript (Gao et al., 2021; Wang et al., 2008). Notably, PurA can also unwind double-stranded DNA, a function critical for transcription and replication (Weber et al., 2016). Our findings suggest that the polyglutamylation of PurA is essential for maintaining colonic homeostasis, which is disrupted in the absence of TTLL6, leading to increased proliferation of colonic epithelial cells. This disruption correlates with enhanced susceptibility to colon carcinogenesis in *Ttll6-*deficient mice and aligns with the observed downregulation of *Ttll6* during colon tumorigenesis in both mice and human CRC. Moreover, mice deficient for *Ttll6* in CEC displayed an elevated tumor progression at the early phase of tumor development. This appears to be associated with the identified mutual dependency of TTLL6 catalyzed polyglutamylation of PurA, the glutamylase itself, and their cellular localization to the nucleus. Additional investigations are needed to clarify the molecular pathways underlying the findings described in this study. It is tempting to speculate that the nuclear localization of PurA and its RNA-binding capacity may regulate gene expression programs that control epithelial proliferation and differentiation, thereby limiting tumor initiation in the colon. Understanding these mechanisms and key molecules may enable the design of novel therapies.

## Acknowledgments

We acknowledge the imaging facility MRI, member of the national infrastructure France-BioImaging (https://ror.org/01y7vt929) supported by the French National Research Agency (ANR-24-INBS-0005 FBI BIOGEN)”. We acknowledge the “Réseau d’Histologie Expérimentale de Montpellier” - RHEM facility supported by SIRIC Montpellier Cancer (Grant INCa_Inserm_DGOS_12553), the European regional development foundation and the occitanian region (FEDER-FSE 2014-2020 Languedoc Roussillon) for processing our animal tissues, histology technics and expertise, as well as the IGMM mouse facility ZEFI. Many thanks to Thierry Gostan and the platform SERANAD for help with the statistical analysis and the support of the funding organizations Cancer Inserm-ITMO Aviesan 2021 (to MH, EM, CJ), ANR (AAPG2021, CILCOL to MH, EM, CJ), FRM (“équipe labellisé”, MH, CJ), La Ligue National contre le Cancer (LNCC2022, to VP) and La Ligue Regional Ligue Régionale contre cancer-Comité de l’Hérault (LRCC (34)2025, to VP). Proteomics analyses were done using the mass spectrometry facility of Marseille Proteomics (marseille-proteomique.univ-amu.fr) supported by IBISA (Infrastructures Biologie Santé et Agronomie), Plateforme Technologique Aix-Marseille, the Cancéropôle PACA, the Provence-Alpes-Côte d’Azur Region, the Institut Paoli-Calmettes, Fonds Européen de Développement Regional (FEDER) and Plan Cancer.

## Material and Methods

### Patient samples

Untreated human samples from colorectal adenomas, adenocarcinoma, liver metastasis and matched normal adjacent colorectal tissues were collected by the surgical oncology and stored in the Biological Resource Center (BRC) of the Institut Curie, France (permission No. A10-024). According to French regulations, patients were informed of research performed with the biological specimens obtained during their treatment and did not express opposition. Disease stage of CRC was based on 7th revised edition of the AJCC Colorectal Cancer (Rocha et al., 2014b). We retrospectively collected clinicopathological and gene expression data for CRC samples from 22 public datasets (Supplementary Table 1). We gathered the transcriptome of 3,333 primary tumors.

### Animal study approval

Mouse experiments were performed in strict accordance with the guidelines of the European Community (directive n°2010/63/EU) and the French National Committee (Project number APAFIS#18685 and #52137) for the care and use of laboratory animals and were approved by the Regional Ethics committee. Statistical analysis was conducted to minimize the number of mice to be used for identifying significant differences.

### Generation of mouse line Ttll6^frt/frt^ and Ttll6 VilinCre-ERT2

The *Ttll6 knock in* mouse line was generated at the Institut Clinique de la Souris – PHENOMIN-ICS (http://www.phenomin.fr). Three embryonic stem cell clones were obtained from the European Mouse Mutant Cell Repository (https://www.eummcr.org/). All clones underwent validation by Southern blot analysis using an internal Neo probe, alongside PCR confirmation of the presence of the 3’ loxP site. Clone EPD0327_6_H04 was further assessed by chromosome spread and Giemsa staining for karyotype analysis, and subsequently microinjected into BALB/cN blastocysts. Male chimeras were obtained, and germline transmission was achieved for the tm1a allele (knock-out first with conditional potential).

*Ttll6^frt/frt^* mice carried a LacZ-neomycin cassette inserted into the *Ttll6* gene, resulting in disruption of its expression (Figure 3A). To generate *Ttll6^flox/flox^* mice, the LacZ-neomycin cassette flanked by FRT sites, was excised by crossing *Ttll6^frt/frt^* mice with mice expressing the FLP recombinase (Kranz et al., 2010). *Ttll6^flox/flox^* mice were then crossed with VilinCre^ERT2^ transgenic mice (el Marjou et al., 2004) to allow epithelial cell specific deletion of exon 9 of *Ttll6* gene (supplementary figure 5). *Ttll6^+/+^*, *Ttll6^frt/frt^* and *Ttll6^flox/flox^*-VilinCre^ERT2^ mice were maintained on C57BL/6 genetic background, and experimental groups including littermates were housed together according to gender and age. Genotyping was done using primers listed in Table 1. Mice used for the in vivo experiments were used at age of 8–10 weeks.

**Table 1.**
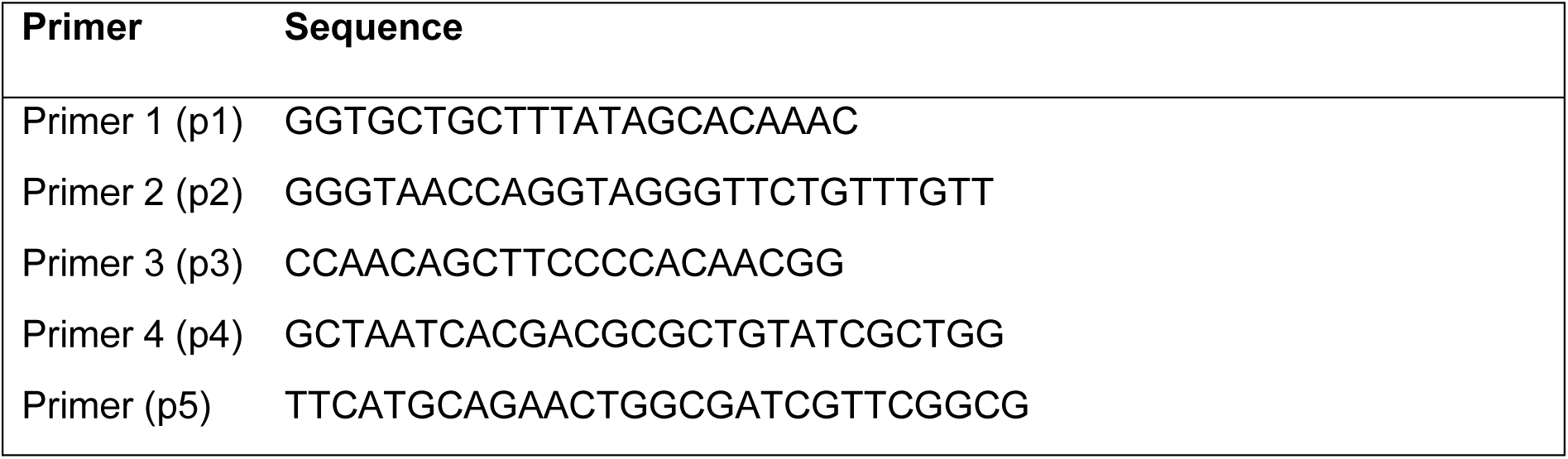
Primers used for genotyping (PCR).

### Genomic DNA extraction and PCR

Genomic DNA was extracted from mice tails, lysate (25mM NaOH, 0.2mM EDTA pH8, mQH2O) for 20min at 95°C followed by neutralization (40mM Tris-HCl pH5). A standard PCR was performed using 2ul of gDNA. Primers used for DNA amplification are listed in Table 1. DNA visualisation were done in 2% agarose gel containing 0.2µg/ml ethidium bromide under UV light.

### RT–qPCR

RT-qPCR was performed with RNA extracted from murine colonic epithelial cells from 8 weeks-old mice using High pure RNA isolation kit (Roche) following the manufacturer’s protocol. RNA was translated to cDNA with SuperscriptIII reverse transcriptase (Invitrogen) using Random Hexamers (Invitrogen). Quantitative RT–PCR was applied under standard conditions using SYBR Green (Roche) on a LightCycler 480 (Roche). The relative mRNA expression levels of each gene were expressed in target gene expression relative to the housekeeping gene *Tbp*. Primers used are listed in the Table 2.

**Table 2.**
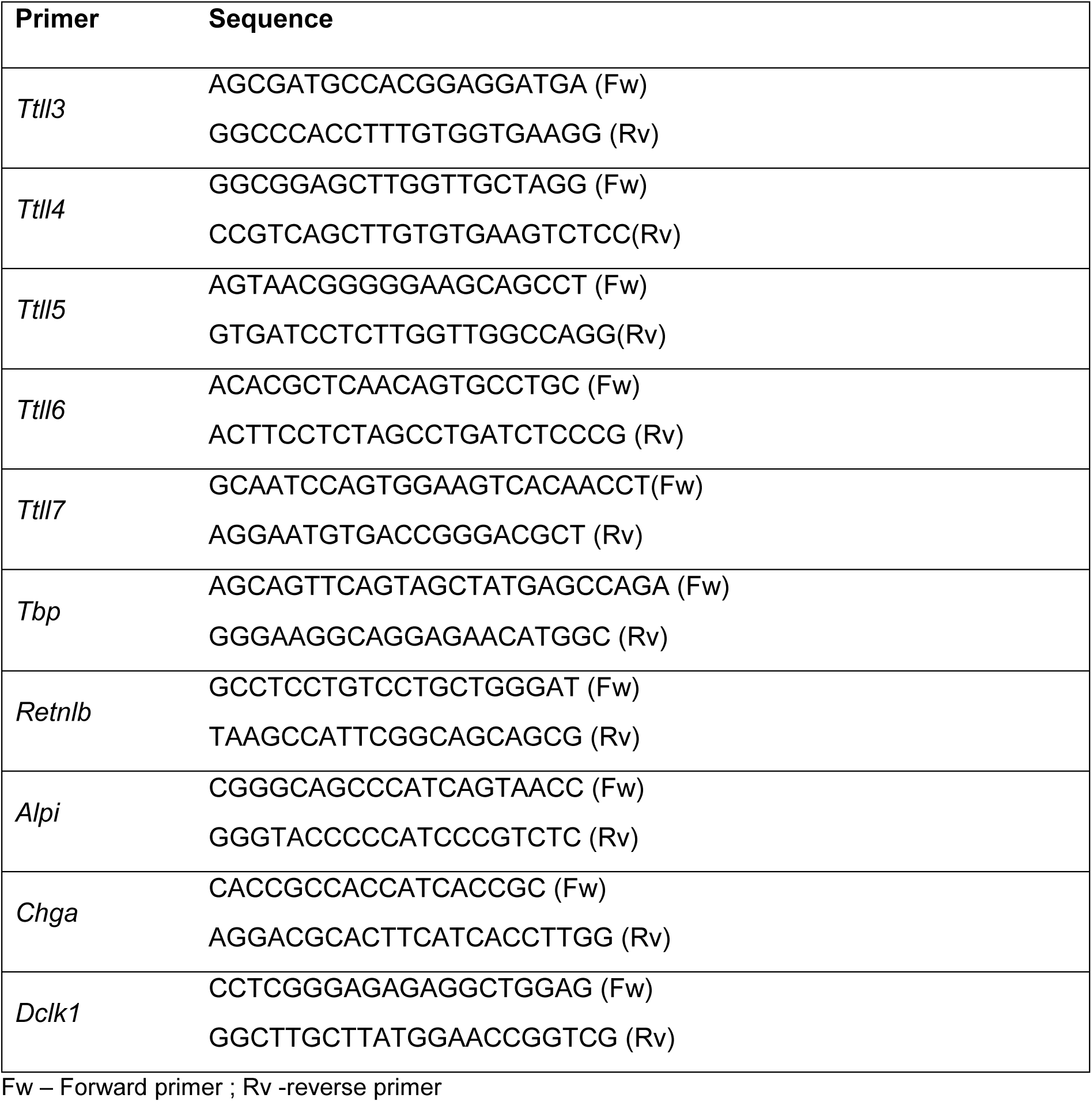
Primers used for RT-qPCR.

### Bulk transcriptome analysis

#### Gene expression

Before analysis, data preprocessing included two successive steps. The first one was to normalize each dataset separately: we applied quantile normalization for the available processed data from non-Affymetrix-based sets and robust multichip average with the nonparametric quantile algorithm for the raw Affymetrix-based datasets. Normalization was performed in R via Bioconductor and associated packages (Taminau et al., 2011). In the second step, hybridization probes were mapped across the different technological platforms represented as previously reported (Bertucci et al., 2014). When multiple probes mapped to the same GeneID, we retained the one with the highest variance in a particular dataset.

Next, we corrected the 22 studies for batch effects via z score normalization (Johnson et al., 2007). Briefly, for each *TTLL6* expression value in each study separately, the value was transformed by subtracting the mean of the gene in that dataset from its standard deviation in the primary colorectal cancer (CRC) samples. Binary values were analyzed on the basis of the quartile of the *TTLL6* expression level in the primary samples. *TTLL6* Low included the first through third quartiles, and *TTLL6* High was defined as the fourth quartile.

#### Statistical analysis

The clinical endpoint of our prognostic analysis was recurrence-free survival (RFS), which was calculated from the date of diagnosis until the date of disease relapse or death. The follow-up time was measured from the date of diagnosis to the date of the last visit for event-free patients. Univariate and multivariate prognostic analyses for RFS were performed via Cox regression analysis (Wald test). The variables tested in the univariate analysis included the sample groups based on *TTLL6* expression, patient age and sex (male, female), location (proximal, distal), pathological Tumour-Node-Metastasis (pTNM) status, differentiation grade, mismatch repair status, and consensus molecular subtype (CMS) classification. Multivariate analysis incorporated all variables with a p value less than 5% in the univariate analysis. Survival was calculated via the Kaplan‒Meier method, and curves were compared via the log-rank test. Survival analysis was performed via the survival package (version 2.43) in R software (version 3.5.2, https://cran.r-project.org).

### Colonic epithelial cell (CEC) extraction

Mice colons were collected, flushed with cold PBS, opened longitudinally and cut in small pieces. Colon pieces were washed three times in ice cold PBS vortexing and pass through 100µm strainer. Colon pieces were incubated in CEC buffer (HBSS, 2% FCS, 5 mM EDTA) at 37°C for 20 minutes on a rotating wheel. Epithelial cells were extracted by vortexing the colon pieces, followed by three washes in ice cold PBS or until colon pieces become transparent. The flow through containing colonic epithelial cells were pellet and snap frozen in liquid nitrogen for further RNA or Protein extraction.

### Colitis-associated carcinogenesis (AOM-DSS)

Mice were injected with tamoxifen intraperitoneally 7 and 5 days before starting the AOM-DSS protocol to induce *Ttll6* deletion. A reboost of a single injection of tamoxifen was done at day 40 to ensure the deletion. For induction of colitis-associated carcinogenesis, male mice from *Ttll6 VilinCre-ERT2* line were intraperitoneally injected with azoxymethane (AOM - Millipore Sigma, 6.5ug/g of body weight). Mice were treated with three cycles of 2% (w/v) dextran sodium sulfate (DSS-TdB Consultancy AB) in the drinking water during 6 days, followed by two weeks of recovering in water. The body weight of the mice was daily followed and in case weight loss was more important than 20% mice were sacrificed. Mice were sacrificed at different time point as defined by the experimental cohort (day 20,40 or 70). Colons were fixed in 10% formalin for 24h and embedded in paraffin for further staining and histological analysis. Tissue sections (4 µm) were stained with hematoxylin-eosin-safran (HES) or processed for immunohistochemistry after deparaffinization and subsequent incubation with the primary antibodies listed in Table 3. Histological grading of AOM/DSS-induced tumours was determined with a blinded genotype according to the described classification of intestinal neoplasia (Washington et al., 2013).

**Table 3.** Antibodies used for Immunohistochemistry (IHC) /Immunofluorescence (IF) and blotting (WB)

| Antigen | Supplier | Clone/Reference | Dilution used | Application |
| --- | --- | --- | --- | --- |
| Ki67 | Abcam | Sp6-ab16667 | 1:200 | IHC |
| BrdU | Biolegend | BU20a-339810 | 1:100 | IHC |
| HES1 | Cell Signalling | D6P2U -11988S | 1:500 | IF |
| CD44v6 | ThermoFisher | MA1-81995 | 1:200 | IF |
| Muc2 | GENETEX | GTX100664 | 1:300 | IHC |
| Vimentin | Cell Signalling | D21H3-5741 | 1:300 | IHC/IF |
| Arl13b | NeuroMab | N295B/66 | 1:1000 | IF |
| PolyE | Adipogen | IN105-AG-25B-0030 | 1:1000/1:2000 | IF/WB |
| GT335 | Adipogen | AG-20B-0020 | 1:1000 | WB |
| GFP | Novus Biologicals | NB100-1770 | 1:500 | IF |
|  | Torrey Pines Biolabs Inc | TP401 | 1:5000 | WB |
| α-tubulin | Sigma | DM1A | 1:1000 | WB |
| HA | Roche | 3F10 | 1:500 | IF |
| PURA | ProteinTech | 67305-1-Ig,clone 1C6G4 | 1:200 | IF |

### Histology and immunohistochemistry

Organs were fixed in 10% neutral buffered formalin (NBF) solution for 24 h. Histological examination was performed on paraffin-embedded sections stained with haematoxylin-eosin-safran and alcian blue staining was also performed. Immunohistochemistry was performed on formalin-fixed and paraffin-embedded tissues cut into 4 µm sections. After blocking of nonspecific binding (with TBS, 10% goat serum, 5% BSA, 5% milk, 0.3% Triton), samples were incubated overnight at 4°C with primary antibody (Table 3). Polymer horseradish peroxidase kits for primary mouse or rabbit antibodies (Vector Laboratories) were used for detection. The revelation was done with 3,30-diaminobenzidine (DAB). For immunofluorescence, tissues samples were incubated overnight at 4°C with primary antibodies and revealed with fluorescent-labelled secondary antibodies (Table 3). DNA was stained using 20 µg/ml of 4’,6’- diamidino-2-phenylindole (DAPI). For the detection of PC, we followed a previously described protocol that allows the detection of PC on paraffin-embedded tissues (Paul et al., 2022).

The RPE1 cells grown on coverslips were co-transfected with wild type HA-PURA or mutated HA-PURA-DD combined with EYFP-TTLL6. Following 24h of transfection, the cells were fixed with -20°C methanol for 5min. After fixation, the cells were washed twice with PBS and used for staining with anti HA and GFP antibodies (Table 3) in PBS, 3% BSA, 0.1% Triton. The coverslips were incubated for 2h at room temperature with the primary antibody dilution, washed once with PBS, 0.1% Triton and incubated additional hour with secondary anti rat antibody conjugated with Alexa-555 (Invitrogen) diluted 1:1000 in PBS, 3% BSA, 0.1% Triton. Following the incubation with secondary antibodies, the cells were stained for 5 min with Hoechst (1mg/ml) diluted 1:1000 in PBS, 0.1% Triton and then washed 4 times with PBS, 0.1% Triton. The coverslips were mounted in Mowiol and then allowed to dry. Immunohistochemistry and immunofluorescence were quantified using the image analysis softwares : QuPath, Image J or NDP view.

### RNAscope in situ hybridization

The RNAscope assay was combined with vimentin immunofluorescence and *Ttll6* probing in paraffine-embedded colon tissue sections with the RNA-Protein Co-Detection Ancillary Kit (Cat# 323180). RNAscope^TM^ Multiplex Fluorescent Reagent Kit v2 (Cat#323110), customized double Z oligonucleotide probes including RNAscope^TM^ Probe-Mm-Ttll6 (Cat#570931), RNAscope^TM^ Positive Control Probe-Mm-Polr2a (Cat#312471), and RNAscope^TM^ Negative Control Probe-DapB (Cat#310043), and opal fluorescent dye including Opal570 (Cat#ASOP570), were purchased from Advanced Cell Diagnostics (ACD, Biotechne). The protocol for colon tissue preparation, pre-treatment, combined immunofluorescence for vimentin, target probe hybridization, hybridization signal amplification, and probe signal staining were performed according to the manufacture instructions.

### BrdU incorporation assay

Eight-week-old mice were intraperitoneally injected with 100 µg/g bromodeoxyuridine (Sigma, B5002-1G). Colons were collected 2 h after injection fixed in 10% formalin for 24h and embedded in paraffin. Proliferating cells were detected by standard immunohistochemistry protocol (described above), with an additional step of DNA denaturation in pre-heated 2M HCl for 30 minutes followed by incubation with sodium borate 0.1M pH 9 for 5 min to preclude DNA strand renaturation. BrdU positive cells were quantified using QuPath software.

### Cell culture

BMDMs were isolated from femurs and tibiae of 6-8w old WT mice. The BM cells were treated with red blood cell lysis buffer (Merk) and suspended in RPMI media with 10% FBS supplemented with 50 ng/ml of M-CSF. Differentiation occurred over 7 days with medium replacement with fresh medium on days 2, 4, and 6.

The protocol to isolate colonic fibroblasts (CF) was modified according to (Paul et al., 2022b). Briefly, colons were isolated from mice (38-42 days old male mice), cut into small pieces, extensively washed, and incubated in HBSS - 2% FCS - 5 mM EDTA buffer at 37°C for 20min with continuous shaking to release CEC. After extensive washing, the tissue was further minced into smaller pieces and digested enzymatically in HBSS - 2% FCS solution containing 62.5 µg/ml Liberase (Sigma) and 40 µg/ml DNase (Sigma). Digestion was performed at 37°C for 50 minutes under continuous shaking with a magnetic stirring device to isolate CF. For the culture, cells were filtered through a 40µm strainer to obtain single cell suspension, and seeded in a flat-bottom 24-well plate. Cells were cultured in DMEM (250mg/ml Strepto-7.5x10^6^U/ml PenicillineG peni/stepto, 10% FCS, 10mM HEPES (Gibco), 2mM sodium pyruvate (Gibco), 50µM beta-mercaptoethanol (Gibco) for two passages and maintained at 37°C in a humidified atmosphere with 5% CO2. The culture medium was replaced every 2 days. The first passage was performed between days 5 and 7 of culture, and the second passage between days 9 and 11. CF were collected between days 15 and 17. HEK293T and RPE1 transfected cells were cultured in Dulbecco’s modified Eagle’s medium/F-12 GlutaMAX-I (Gibco) supplemented with 10% heat-inactivated fetal bovine serum (Gibco) and antibiotics (penicillin/streptomycin) (Gibco). For all cell lines, transfection of expression plasmids was performed with JetPEI transfection reagent (Polyplus) according to manufacturer’s guidelines.

### Transfection

HEK293T cells were co-transfected with either pEGFP_C1-PURA or pEGFP_C1-PURB and pEYFP-TTLL4_C639 or pEYFP-TTLL6_N513 plasmid using JetPEI transfection reagent (Polyplus) according to manufacturer’s guidelines. The Ttll4 and Ttll6 expressing plasmids have been previously described (van Dijk et al., 2007). Site-specific mutagenesis of the *purA* gene to generate pEGFP_C1-PURA-DDD was introduced by PCR prior to transfection.

RPE1 cells were transfected with pRK5-HA-PURA or pRK5-HA-PURA-DDD either alone or in combination with pEYFP-TTLL6_N513 plasmid.

### Tumoroid culture

Tumoroids were generated by isolating colonic tumour lesions from AOM/DSS treated mice, cut in small pieces, washed three times in PBS and subsequent enzymatically dissociation with TrypLE express (Gibco) and DNase at 37°C for 30min. The tumoral cells were cultured in reduced growth factor Matrigel (Corning) in Advanced DMEM/F12 supplemented with Glutamax, HEPES, B27, N2, N-acetyl cysteine, mouse EGF and Noggin. The medium was renewed every 3-4 days and cells were passed when confluent. Tumoroids were collected between second to fourth passage for RNA extraction.

### Immunoblot

Following 24h transfection, protein samples were prepared in 1x Laemmli buffer and 7-10 µg of total proteins from each sample was separated by SDS-polyacrylamide gel electrophoresis (Magiera and Janke, 2013) and transferred onto nitrocellulose membranes. The membranes were incubated with skim milk to block non-specific binding before incubation with specific primary and secondary antibodies, respectively. The following primary antibodies (Table 3) were used: GT335 (AdipoGen), PolyE (AdipoGen), anti HA (Roche) and anti GFP (Torrey Pines Biolabs). The intensity of the protein bands was determined by using a CCD camera.

### Immunoprecipitation

Colonic epithelial cells from distal and transverse segment from eight-weeks old mice were collected and snap freeze. Cells were ground to powder under liquid nitrogen. The powder was resuspended in protein lysis buffer (50mM Hepes pH7.3, 10%glycerol,150mM NaCl,1%triton-X-100, 0.5mM DTT, mQH2O) containing proteases inhibitors and phosphatases. The samples were let on ice for 30min and centrifuged at 16000g for 30min at 4°C. The supernatant was incubated overnight at 4°C with Dynabeads protein G-coupled magnetic beads (Invitrogen) coated with anti-polyE antibody (AdipoGen). The beads were then washed three times with PBS containing 0.01% Tween20 and beads were eluted in 30ul of 2xLaemlli containing 200mM DTT and boiled for 5 min at 95°C. The bound proteins were analysed by pierce silver stain, immunoblot and analysed by mass-spectrometry.

### Mass spectrometry and data analysis

Immunoprecipitated samples were loaded on NuPAGE™ 4–12% Bis–tris acrylamide gels according to the manufacturer’s instructions (Life Technologies). Running of protein was stopped as soon as proteins enter the top of the gel and stacked in a single band. Protein containing bands were stained with Imperial Blue (Pierce), cut from the gel and digested with high sequencing grade trypsin (Promega) before mass spectrometry analysis. Gel pieces were washed and destained using few steps of 100mM NH_4_HCO_3_. Destained gel pieces were shrunk with 100 mM ammonium bicarbonate in 50% acetonitrile and dried at RT. Protein spots were then rehydrated using 10mM DTT in 25 mM ammonium bicarbonate pH 8.0 for 45 min at 56°C. This solution was replaced by 55 mM iodoacetamide in 25 mM ammonium bicarbonate pH 8.0 and the gel pieces were incubated for 30 min at room temperature in the dark. They were then washed twice in 25 mM ammonium bicarbonate and finally shrunk by incubation for 5 min with 25 mM ammonium bicarbonate in 50% acetonitrile. The resulting alkylated gel pieces were dried at room temperature. The dried gel pieces were reswollen by incubation in 25 mM ammonium bicarbonate pH 8.0 supplemented with 12.5 ng/µl trypsin (Promega) for 1h at 4°C and then incubated overnight at 37°C. Peptides were harvested by collecting the initial digestion solution and carrying out two extractions; first in 5% formic acid and then in 5% formic acid in 60% acetonitrile. Pooled extracts were dried down in a centrifugal vacuum system. Samples were reconstituted in 0.1% TFA 4% acetonitrile before mass spectrometry using an Orbitrap Fusion Lumos Tribrid Mass Spectrometer online with an Vanquish Neo chromatography system (ThermoFisher Scientific, San Jose, CA). Peptides were separated at 40°C using a two steps linear gradient (4–20% acetonitrile/H2O; 0.1% formic acid for 110 min and 20-32% acetonitrile/H2O; 0.1% formic acid for 10 min). An EASY-Spray nanosource was used for peptide ionization (1,900 V, 275°C). MS was conducted using a data-independent acquisition mode (DIA). Full MS scans were acquired in the range of m/z 375–1,500 at a resolution of 120,000 at m/z 200 and the automatic gain control (AGC) was set at 4.0 × 10E5 with a 50 ms maximum injection time. MS2 spectra were acquired in the Orbitrap with a resolution of 30,000, in the mass range of 200-1800 m/z after isolation of parent ion in the quadrupole and fragmentation in the HCD cell under collision Energy of 30%. DIA parent ion range was from 400 to 1000 m/z divided into 40 windows 16 Da wide and from 1000 to 1500 m/z divided into 10 windows 50 Da wide.

For protein identification and quantification, relative intensity-based label-free quantification (LFQ) was processed using the DIA-NN 1.8 algorithm (Demichev, et al., 2020). Raw files were searched against the *Mus Musculus* database extracted from UniProt (UP000000589, date 2025-03-13 containing 54727 entries) implemented with a contaminant database (Frankenfield, et al., 2022). The following parameters were used for searches: (i) trypsin allowing cleavage before proline; (ii) one missed cleavage was allowed; (iii) cysteine carbamidomethylation (+57.02146) as a fixed modification and methionine oxidation (+15.99491) and N-terminal acetylation (+42.0106) as variable modifications; (iv) a maximum of 1 variable modification per peptide allowed; and (v) minimum peptide length was 7 amino acids and a maximum of 30 amino acids. The match between runs option was enabled. The precursor false discovery was set to 1%. DIA-NN parameters were set on double-pass mode for Neural Network classifier, Robust LC High precision for quantification strategy and RT-dependent mode for Cross-run normalization. Library was generated using Smart profiling set up. MS1 and MS2 mass accuracy was automatically calculated for precursor charge fixed between 2 and 4.

Main output file from DIA-NN was further filtered at 1% FDR and LFQ intensity was calculated using our DIAgui package at 1% q-value (https://github.com/marseille-proteomique/DIAgui (Gerault et al., 2024). The statistical analysis was done with Perseus software (version 1.6.15.0), where proteins were pre-filtered to remove contaminants (Tyanova and Cox, 2018). Relative quantification was calculated using fold changes of LFQ intensity between both conditions (WT, KO TTL6). To determine whether a given detected protein was specifically differential, a two-sample *t*-test was done using permutation-based FDR-controlled employing 250 permutations. The mass spectrometry proteomics data have been deposited to the ProteomeXchange Consortium via the PRIDE (Perez-Riverol et al., 2025) partner repository with the dataset identifier PXD070756.

### Microscopy and imaging

Histological slides were scanned using Nanozoomer 2.0 HT scanner with a 40x objective and visualized with NDP.view2 viewing software (Hamamatsu). Fluorescent Images were acquired using an Andor Dragonfly Spinning Disk Confocal inverted microscope using a 40X oil immersion objective. 26 Z-stack sections were captured at 0.2 µm intervals over a 5 µm range. RNAscope fluorescent images were acquired using the upright Leica Thunder imager with LAS X software.

### Statistical analysis

All statistical analysis of quantifications for the mouse study were performed in GraphPad Prism version 10 using a 2-tailed unpaired student t-test when data passed the normality tests (Kolmogorov-Smirnov (KS) normality test, D’Agostino–Pearson omnibus normality test, Shapiro–Wilk normality test). If values were not normally distributed, a nonparametric Mann–Whitney U test was used. For the Ki67 and BrdU statistical analysis quantification using ordinary two-way ANOVA-Šídák’s multiple comparisons test.

