## Supplemental data for "TTLL6-mediated Polyglutamylation of PurA Maintains Colonic Crypt Integrity"

*Supplementary data*

### ***Supplementary figure 1***

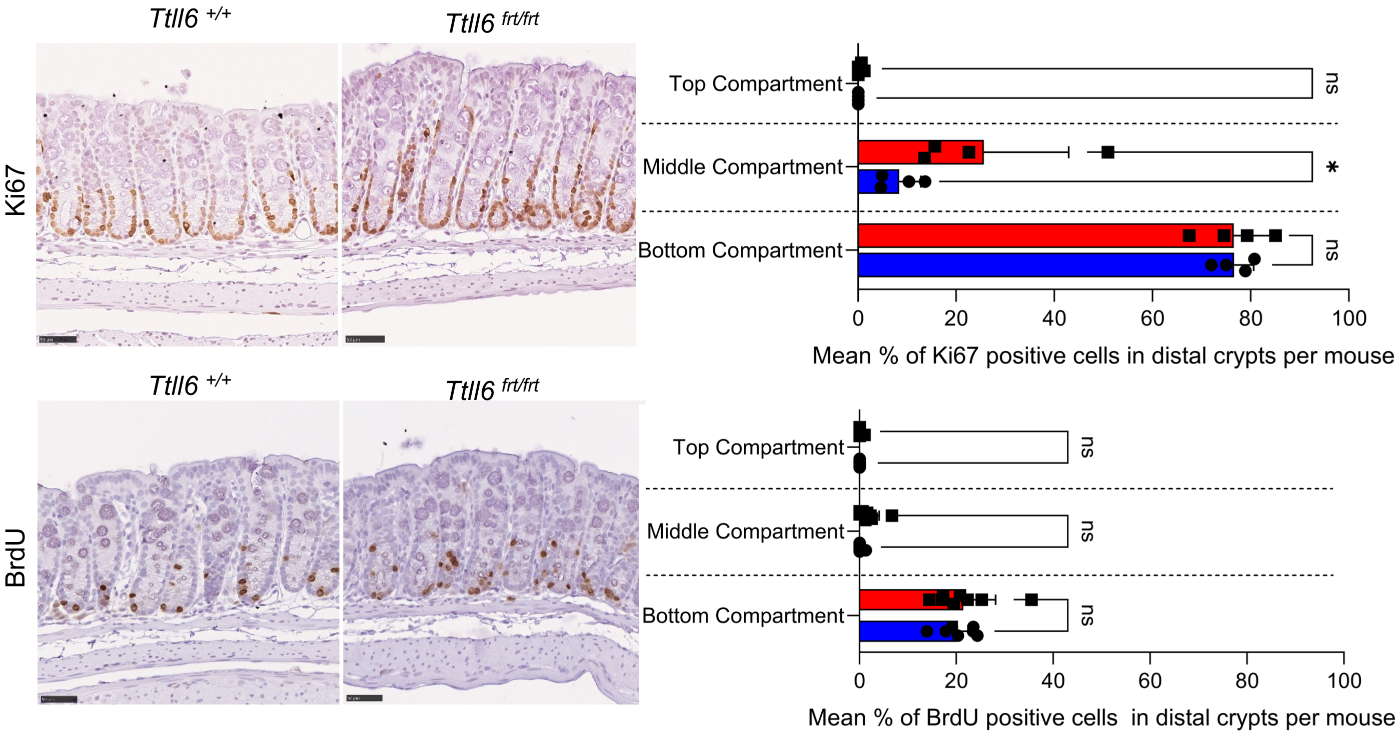


***Supplementary figure 1.*** Top panel :Representative images of immunohistochemistry of Ki67 staining in distal segment (n=4 for each group *Ttll6*$^{+/+}$ (blue) and *Ttll6*$^{frt/frt}$ (red)) with the respective quantification per cell compartment in %. Bottom panel : Representative images of immunohistochemistry of BrdU pulse 2h in distal segment ((n=6 for *Ttll6*$^{+/+}$ (blue) and n=7 for *Ttll6*$^{frt/frt}$ (red)) with the respective quantification per cell compartment in %.

Per mouse 10-40 open crypts were quantified per segment (B,C,E,F). Scale bar 50um.

Statistical analysis : ns p>0.05, * $p < 0.05$, **$p < 0.01$, ***$p < 0.001$, ****$p < 10-4$ by ordinary two-way ANOVA-Šídák's multiple comparisons test

*Supplementary Figure 2*

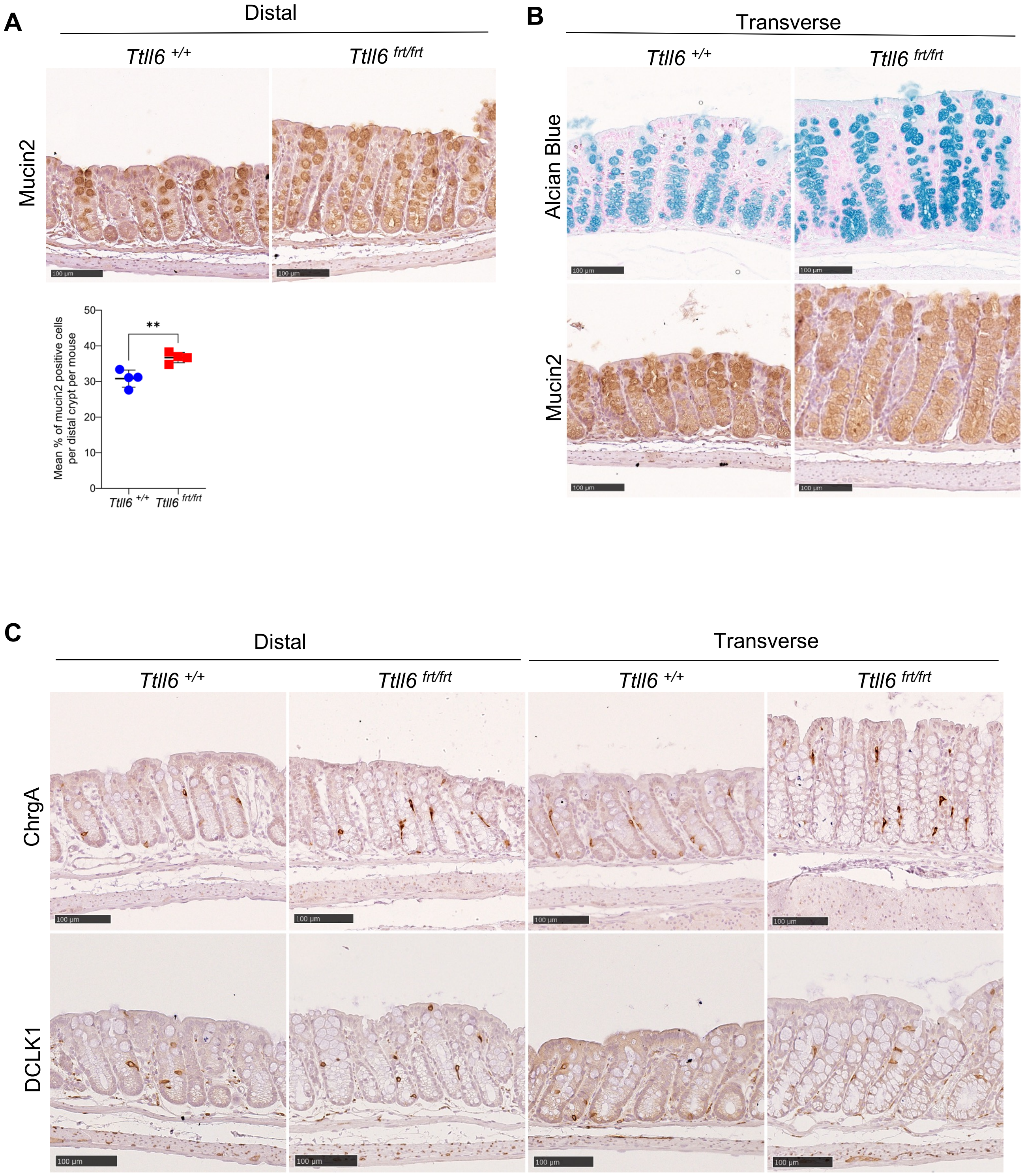


**Supplementary figure 2. (A)** Representative images of immunohistochemical analysis for mucin2 positive cells in distal segments (n=4 for each group *Ttll6*$^{+/+}$ (blue) and *Ttll6*$^{frt/frt}$ (red)) , visualized using mucin 2 and respective quantification of the percentage of mucin2 positive cells per crypts per mouse. **(B)** Increase number mucinous positive cells in *Ttll6*-deficient mice (*Ttll6*$^{frt/frt}$) in transverse segment. Representative images of immunohistochemistry for mucin cell (eg. goblet cells) markers (Alcian blue and Mucin2) in transverse segment . **(B)** There are no differences in enteroendocrine and tuft cells. Representative images of immunohistochemistry for different types of differentiated cells as: enteroendocrine cells using marker chromogranin A (ChrgA) and tuft cells using the marker DCLK1 in distal and transverse segments. Scale bar 100um. Statistical analysis: p>0.05, * $p < 0.05$, ** $p < 0.01$, *** $p < 0.001$, **** $p < 10-4$ by two-tailed unpaired t-test.

### *Supplementary Figure 3*

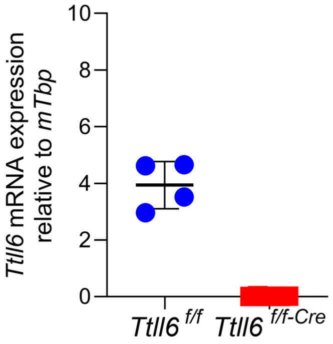


***Supplementary figure 3.****Ttll6* transcript levels in colonic epithelial cells from *Ttll6flx/flx-vilincreERT2* mice, 4 weeks upon tamoxifen administration. Blue dot represent transcript levels of *wild-type (Ttll6*$^{f/f}$, n=4*)* , and red dot of *Ttll6*-deficient *(Ttll6*$^{f/f\text{-}Cre}$*,* n=6*)* mice*. Ttll6* transcript levels were normalized to mouse TATA-binding protein (*Tbp)*

### *Supplementary figure 4*

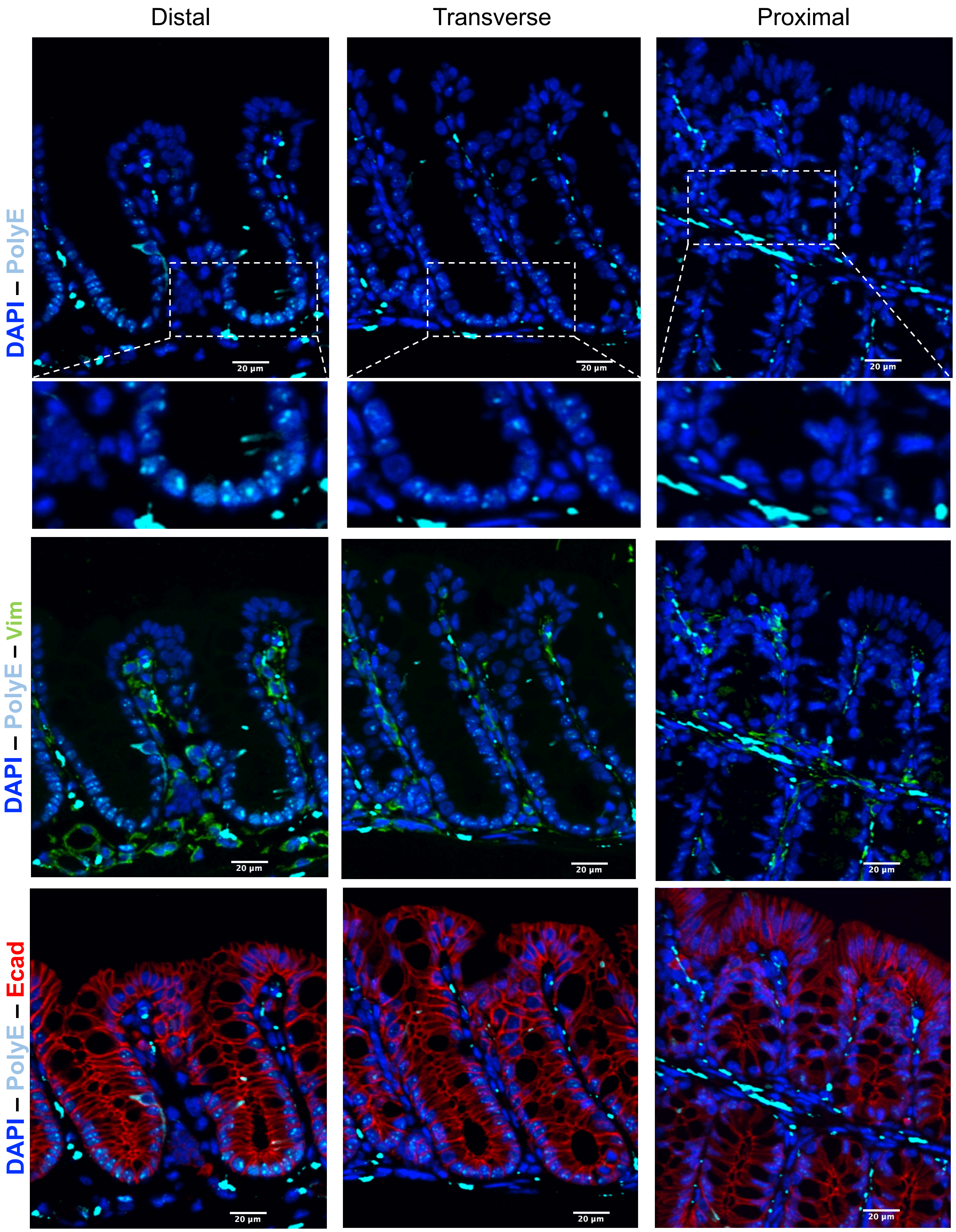


***Supplementary figure 4.*** **In the colon, epithelial cell polyglutamylation status varies between distal, transverse and proximal segments.** In *Ttll6*$^{+/+}$ colon, epithelial cells were labeled with E-Cadherin (in red), fibroblasts with vimentin (in green) and nuclei with DAPI (in blue). Nuclear punctuated staining for polyglutamylation (in cyan) is detectable in epithelial cells with a decreasing gradient from distal, to transverse and proximal parts. Scale bar represent 20µm. A 2x magnification of the DAPI-PolyE staining is shown below the corresponding images.

### *Supplementary figure 5*

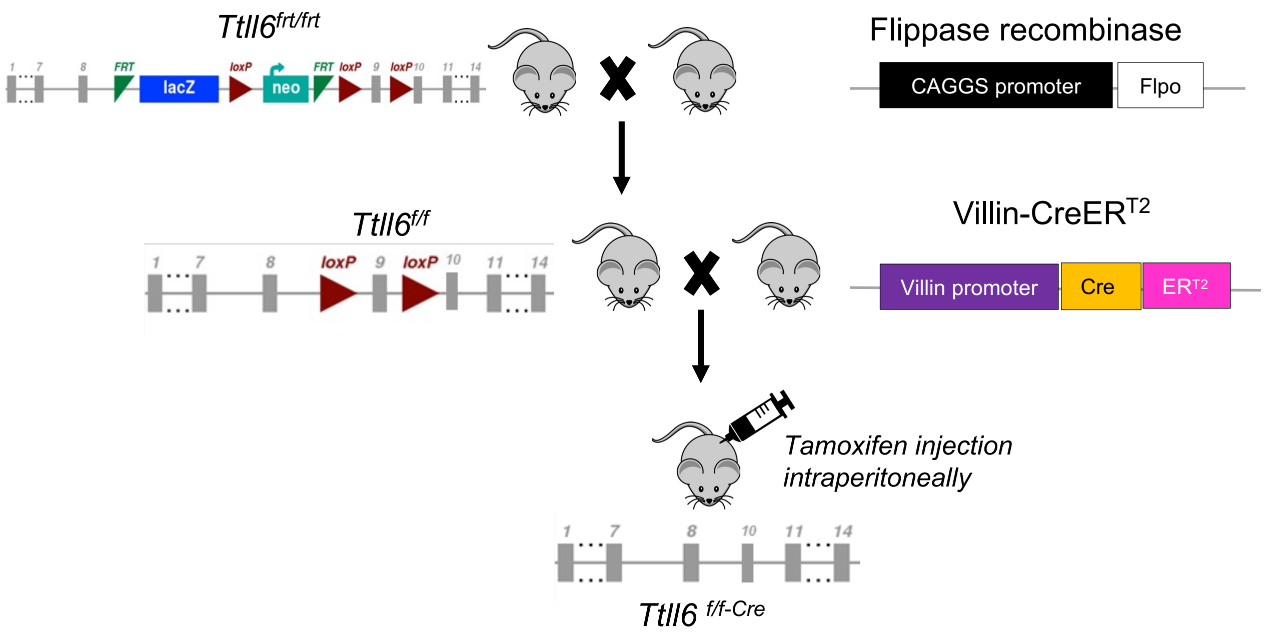


**Supplementary figure 5 . Generation of *Ttll6* VilinCre-ERT2 strain.** *Ttll6*$^{frt/frt}$ mice carry a LacZ-neomycin cassette inserted into the *Ttll6* gene, disrupting its expression. To generate *Ttll6*$^{flox/flox}$ mice, the LacZ-neomycin cassette flanked by FRT sites, was excised by crossing *Ttll6*$^{frt/frt}$ mice with mice ubiquitously expressing the FLP recombinase. The resulting *Ttll6 flox/flox* (*Ttll6*$^{f/f}$) mice were then crossed with VillinCre-ERT2 transgenic mice. The Villin promoter is tissue-specific, primarily driving gene expression in the epithelial cells of small intestine and colon. Cre recombinase is fused to a mutated ligand-binding domain of the estrogen receptor (ERT2). In the absence of tamoxifen, the estrogen receptor domain sequesters Cre in the cytoplasm. Upon tamoxifen administration, Cre-ERT2 translocates to the nucleus, activating Cre recombinase specifically in cells expressing the Villin promoter. This strategy enables epithelial cell-specific deletion of exon 9 of the *Ttll6* gene.

### *Supplementary figure 6*

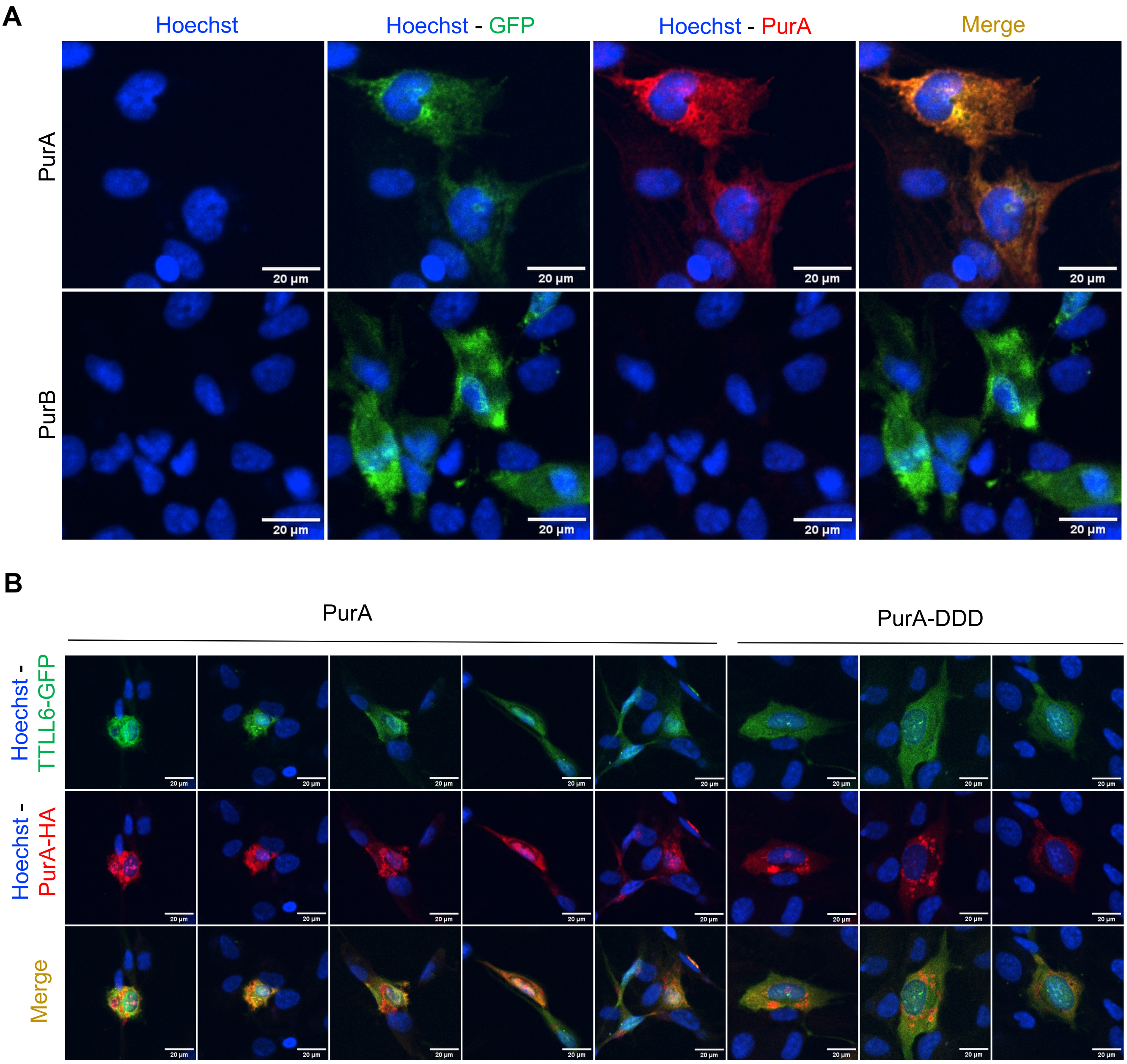


**Supplementary figure 6. (A)** *PurA antibody validation in RPE cells transfected with PurA or PurB.* Representative images of immunofluorescence analysis of RPE cells transfected with PURA-EGFP (top panel, green) or PurB-EGFP (bottom panel, green) and staining with PurA-EGFP, revealing specific detection of PurA and not PurB. **(B)** Images of immunofluorescence analysis of retinal pigment epithelial (RPE) cells co-transfected with TTLL6-EYFP and PurA-HA or PurA-DDD-HA (mutated form). Upon PurA and TTLL6 co-transfection, both proteins are present in nucleus as well as cytoplasm (top panel). In contrast, in cells co-transfected with TTLL6 and PurA mutated (PurA DDD), only TTLL6 is detected in nucleus and cytoplasm, whereas PurA DDD is located mainly in cytoplasm (bottom panel). Scale bar 20um.

***Supplementary figure 7***

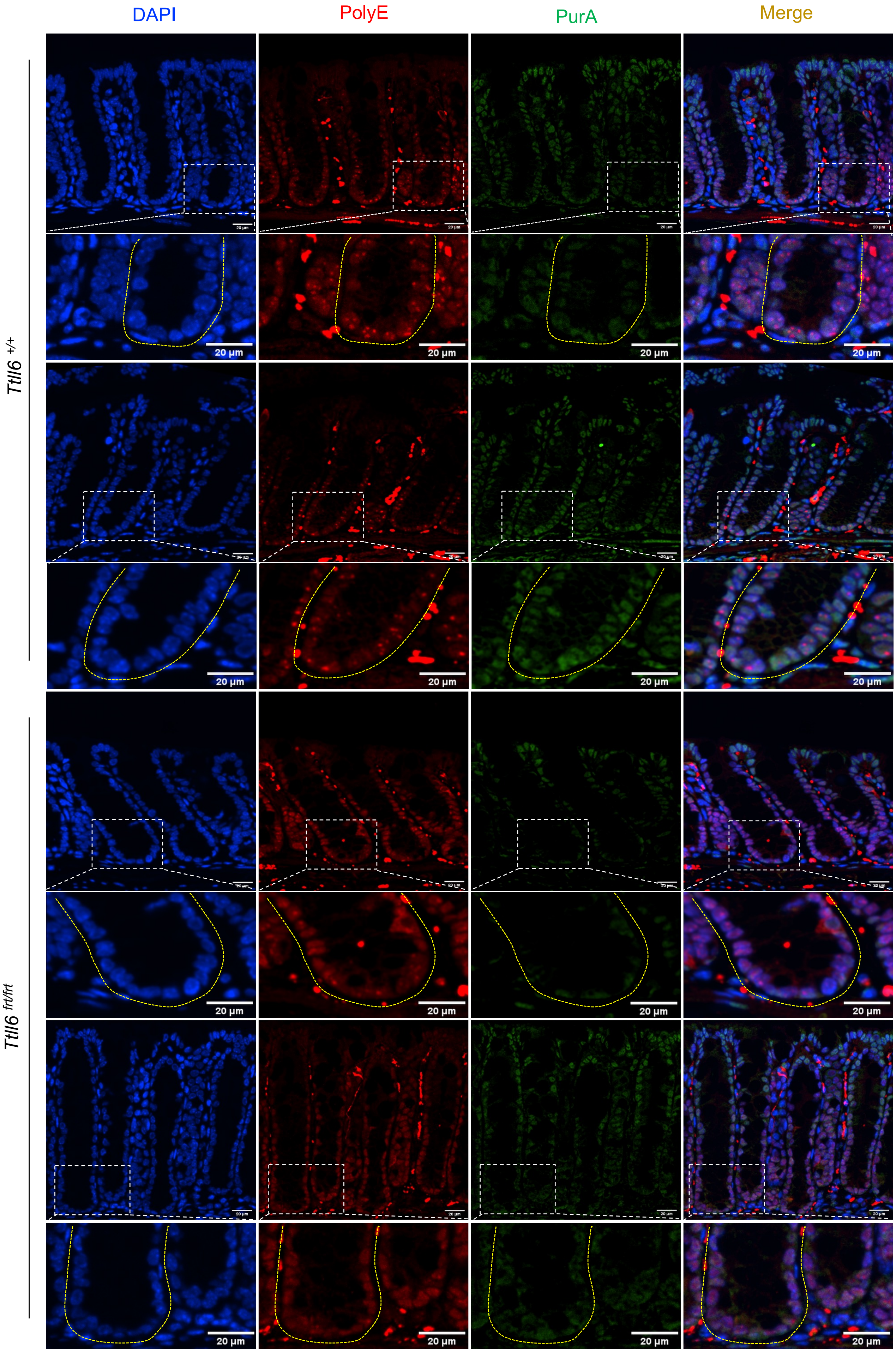

**Supplementary figure 7.** Representative images of immunostaining for polyglutamylation (polyE in red) and purA (in green) of mouse colon sections. In *Ttll6* $^{+/+}$ mice, polyE staining is detectable in bottom and middle crypt compartments with accumulation in nuclear foci. PurA is present in all nuclei of CEC in the crypt. In *Ttll6* $^{frt/frt}$ mice, polyE staining is reduced and not detectable in nuclear foci; PurA staining in nuclei is reduced in bottom and middle crypt compartment, and in the top compartment not all nuclei reveal PurA staining. Images are representative for samples collected from 3 control and 3 *Ttll6*-deficient mice. Scale bar 20um.

**Supplementary Table 1.** Description of the public colon data sets

| Reference | Source of data | Technological platform | N° of probe sets/genes | Primary (N) |
|---|---|---|---|---|
| Expression Project for Oncology (expO), 2005 | GEO database, GSE2109 | Affymetrix, array U133 Plus 2.0 | 54K | 300 |
| Jorissen et al., Clin Cancer Res 2008 | GEO database, GSE13294; GSE13067 | Affymetrix, array U133 Plus 2.0 | 54K | 229 |
| Jorissen et al., Clin Cancer Res 2009 | GEO database, GSE14333 | Affymetrix, array U133 Plus 2.0 | 54K | 251 |
| Gyorffy et al., PLoS One 2009 | GEO database, GSE4183 | Affymetrix, array U133 Plus 2.0 | 54K | 15 |
| Smith et al., Gastroenterology 2010 | GEO database, GSE17538 | Affymetrix, array U133 Plus 2.0 | 54K | 232 |
| Matsuyama et al., Int J Cancer 2010 | GEO database, GSE18105 | Affymetrix, array U133 Plus 2.0 | 54K | 94 |
| Hong et al., Clin Exp Metastasis 2010 | GEO database, GSE9348 | Affymetrix, array U133 Plus 2.0 | 54K | 70 |
| Skrzypczak et al., PLoS One 2010 | GEO database, GSE20916 | Affymetrix, array U133 Plus 2.0 | 54K | 111 |
| de Sousa et al., Cell Stem Cell 2011 | GEO database, GSE33113 | Affymetrix, array U133 Plus 2.0 | 54K | 90 |
| Kennedy et al., J Clin Oncol 2011 | Array-Express database, E-MTAB-863 ; E-MTAB-864 | Affymetrix custom array, ADXCRCG2a520319 | 62K | 359 |
| Sveen et al., Genome Med 2011 | GEO database, GSE24551 | Affymetrix, Human Exon 1.0 ST array | 22K | 160 |
| Uddin et al., Am J Pathol 2011 | GEO database, GSE23878 | Affymetrix, array U133 Plus 2.0 | 54K | 35 |
| Gröne et al., Int J Colorectal Dis 2011 | GEO database, GSE18088 | Affymetrix, array U133 Plus 2.0 | 54K | 53 |
| Vilar et al., Cancer Res 2011 | GEO database, GSE266882 | Affymetrix, array U133 A | 22K | 176 |
| Laibe et al., OMICS 2012 | GEO database, GSE37892 | Affymetrix, array U133 Plus 2.0 | 54K | 130 |
| TCGA, COAD | TCGA portal, https://tcga-data.nci.nih.gov | Illumina, RNA sequencing V2 | 25K | 459 |
| Schlicker et al., BMC Med Genomics 2012 | GEO database, GSE35896 | Affymetrix, array U133 Plus 2.0 | 54K | 62 |
| Marisa et al., PLoS Med 2013 | GEO database, GSE39582 | Affymetrix, array U133 Plus 2.0 | 54K | 454 |
| Isella et al., Nat Commun 2017 | GEO database, GSE73255 | Illumina, HT12 | 45K | 3 |
| Institut Paoli-Calmette | Array-Express database, E-MTAB-11754 | Affymetrix, array U133 Plus 2.0 | 54K | 50 |
| **Total** | | | | 3333 |

**Supplementary Table 2.** Uni- and Multivariate analyses of the association of TTLL6$^{high}$ expression with recurrence free-survival

| RFS | | Univariate N | Univariate HR [95%CI] | Univariate p-value | Multivariate N | Multivariate HR [95%CI] | Multivariate p-value |
|---|---|---|---|---|---|---|---|
| Age_Diag | | 1817 | 1.00 [1.00-1.01] | 0.224 | | | |
| Age_Diag50 | >50 vs. <=50 | 1817 | 1.14 [0.85-1.54] | 0.386 | | | |
| Sex | male vs. female | 1821 | 1.08 [0.91-1.29] | 0.37 | | | |
| Location_2K | proximal vs. distal | 1553 | 0.97 [0.81-1.17] | 0.777 | | | |
| pStage | 2 | 1688 | 5.24 [2.33-11.8] | **1.39E-09** | 1340 | 4.44 [1.64-11.99] | **3.28E-03** |
| | 3 | | 8.39 [3.71-19.0] | | 1340 | 6.95 [2.56-18.88] | **1.42E-04** |
| Grade | 2 | 540 | 1.01 [0.54-1.87] | 0.166 | | | |
| | 3 | | 1.50 [0.74-3.01] | | | | |
| MSI_Banerjea2004 | MSS- vs. MSI-like | 1981 | 1.14 [0.93-1.40] | 0.200 | | | |
| CMS | CMS2 | 1717 | 0.92 [0.71-1.20] | **4.90E-03** | 1340 | 0.87 [0.62- 1.24] | 0.449 |
| | CMS3 | | 1.09 [0.80-1.47] | | 1340 | 1.11 [0.75- 1.63] | 0.613 |
| | CMS4 | | 1.36 [1.05-1.76] | | 1340 | 0.97 [0.67- 1.42] | 0.887 |
| TTLL6_Q4 | Q4 vs. Q1-Q3 | 1844 | 0.68 [0.54-0.85] | **6.24E-04** | 1340 | 0.74 [0.55- 1.00] | **4.74E-02** |

**Supplementary Table 3.** Polyglutamylated proteins enriched in *wild-type* colonic epithelial cells

| Protein ID | Protein name | | Gene |
|---|---|---|---|
| A0A140T8N3 | A0A140T8N3_MOUSE | Immunoglobulin kappa chain variable 13-84 (Fragment) | Igkv13-84 |
| Q3UBX0 | TM109_MOUSE | Voltage-gated monoatomic cation channel TMEM109 | Tmem109 |
| P01867 | IGG2B_MOUSE | Immunoglobulin heavy constant gamma 2B | Ighg2b |
| O54879 | HMGB3_MOUSE | High mobility group protein B3 | Hmgb3 |
| Q8VCZ2 | TM45B_MOUSE | Transmembrane protein 45B | Tmem45b |
| P01872 | IGHM_MOUSE | Immunoglobulin heavy constant mu | Ighm |
| Q91Z31 | PTBP2_MOUSE | Polypyrimidine tract-binding protein 2 | Ptbp2 |
| P42669 | PURA_MOUSE | Transcriptional activator protein Pur-alpha | Pura |
| Q62448 | IF4G2_MOUSE | Eukaryotic translation initiation factor 4 gamma 2 | Eif4g2 |
| O35295 | PURB_MOUSE | Transcriptional regulator protein Pur-beta | Purb |

**Supplementary Table 4.** Polyglutamylated proteins enriched in *wild-type* compared to Ttll6 deficient colon epithelial cells

| Protein ID | Protein name | | Genes |
|---|---|---|---|
| Q9Z2C8 | YBOX2_MOUSE | Y-box-binding protein 2 | Ybx2 |
| P70372 | ELAV1_MOUSE | ELAV-like protein 1 | Elavl1 |
| Q3UYV9 | NCBP1_MOUSE | Nuclear cap-binding protein subunit 1 | Ncbp1 |
| Q9JKB3 | YBOX3_MOUSE | Y-box-binding protein 3 | Ybx3 |
| Q6PHQ9 | Q6PHQ9_MOUSE | Polyadenylate-binding protein | Pabpc4 |
| A2RSY6 | TRM1L_MOUSE | TRMT1-like protein | Trmt1l |
| P23249 | MOV10_MOUSE | Putative helicase MOV-10 | Mov10 |
| P29341 | PABP1_MOUSE | Polyadenylate-binding protein 1 | Pabpc1 |
| Q920Q6 | MSI2H_MOUSE | RNA-binding protein Musashi homolog 2 | Msi2 |
| P01029 | CO4B_MOUSE | Complement C4-B | C4b |
| P62960 | YBOX1_MOUSE | Y-box-binding protein 1 | Ybx1 |
| Q8VC31 | CCDC9_MOUSE | Coiled-coil domain-containing protein 9 | Ccdc9 |
| Q8BI72 | CARF_MOUSE | CDKN2A-interacting protein | Cdkn2aip |
| Q3ULL6 | Q3ULL6_MOUSE | UPF3 regulator of nonsense transcripts homolog B (yeast) | Upf3b |
| Q9EPU0 | RENT1_MOUSE | Regulator of nonsense transcripts 1 | Upf1 |
| Q8CJG0 | AGO2_MOUSE | Protein argonaute-2 | Ago2 |
| Q91Z31 | PTBP2_MOUSE | Polypyrimidine tract-binding protein 2 | Ptbp2 |
| P28659 | CELF1_MOUSE | CUGBP Elav-like family member 1 | Celf1 |
| Q9CQ49 | NCBP2_MOUSE | Nuclear cap-binding protein subunit 2 | Ncbp2 |
| Q8CJG1 | AGO1_MOUSE | Protein argonaute-1 | Ago1 |
| P52479 | UBP10_MOUSE | Ubiquitin carboxyl-terminal hydrolase 10 | Usp10 |
| P63073 | IF4E_MOUSE | Eukaryotic translation initiation factor 4E | Eif4e |
| Q80U78 | PUM1_MOUSE | Pumilio homolog 1 | Pum1 |
| A2AJ72;Q3TIX6 | A2AJ72_MOUSE;Q3TIX6_MOUSE | Far upstream element (FUSE) binding protein 3 | Fubp3 |
| O35295 | PURB_MOUSE | Transcriptional regulator protein Pur-beta | Purb |
| Q8BGD9 | IF4B_MOUSE | Eukaryotic translation initiation factor 4B | Eif4b |
| P61979 | HNRPK_MOUSE | Heterogeneous nuclear ribonucleoprotein K | Hnrnpk |
| Q8VDP4 | CCAR2_MOUSE | Cell cycle and apoptosis regulator protein 2 | Ccar2 |
| Q8C5Q4 | GRSF1_MOUSE | G-rich sequence factor 1 | Grsf1 |
| Q99K48 | NONO_MOUSE | Non-POU domain-containing octamer-binding protein | Nono |
| Q8BWW4 | LARP4_MOUSE | La-related protein 4 | Larp4 |
| Q61990 | PCBP2_MOUSE | Poly(rC)-binding protein 2 | Pcbp2 |
| Q62189 | SNRPA_MOUSE | U1 small nuclear ribonucleoprotein A | Snrpa |
| Q8BYK6 | YTHD3_MOUSE | YTH domain-containing family protein 3 | Ythdf3 |
| Q6NZJ6 | IF4G1_MOUSE | Eukaryotic translation initiation factor 4 gamma 1 | Eif4g1 |
| Q921F2 | TADBP_MOUSE | TAR DNA-binding protein 43 | Tardbp |
| P97855 | G3BP1_MOUSE | Ras GTPase-activating protein-binding protein 1 | G3bp1 |
| A2AT37 | RENT2_MOUSE | Regulator of nonsense transcripts 2 | Upf2 |
| Q6ZQ58 | LARP1_MOUSE | La-related protein 1 | Larp1 |
| Q7TMK9 | HNRPQ_MOUSE | Heterogeneous nuclear ribonucleoprotein Q | Syncrip |
| Q91WT8 | RBM47_MOUSE | RNA-binding protein 47 | Rbm47 |
| Q9WTX2 | PRKRA_MOUSE | Interferon-inducible double-stranded RNA-dependent protein kinase activator A | Prkra |
| Q9DBR1 | XRN2_MOUSE | 5'-3' exoribonuclease 2 | Xrn2 |
